# The global distribution and environmental drivers of the soil antibiotic resistome

**DOI:** 10.1101/2022.07.11.499543

**Authors:** Manuel Delgado-Baquerizo, Hang-Wei Hu, Fernando T. Maestre, Carlos A. Guerra, Nico Eisenhauer, David J. Eldridge, Yong-Guan Zhu, Qing-Lin Chen, Pankaj Trivedi, Shuai Du, Thulani P. Makhalanyane, Jay P. Verma, Beatriz Gozalo, Victoria Ochoa, Sergio Asensio, Ling Wang, Eli Zaady, Javier G. Illán, Christina Siebe, Tine Grebenc, Xiaobing Zhou, Yu-Rong Liu, Adebola R. Bamigboye, José L. Blanco-Pastor, Jorge Duran, Alexandra Rodríguez, Steven Mamet, Fernando Alfaro, Sebastian Abades, Alberto L. Teixido, Gabriel F. Peñaloza-Bojacá, Marco Molina-Montenegro, Cristian Torres-Díaz, Cecilia Perez, Antonio Gallardo, Laura García-Velázquez, Patrick E. Hayes, Sigrid Neuhauser, Ji-Zheng He

## Abstract

**Background:** Little is known about the global distribution and environmental drivers of key microbial functional traits such as antibiotic resistance genes (ARGs). Soils are one of Earth’s largest reservoirs of ARGs, which are integral for soil microbial competition, and have potential implications for plant and human health. Yet, their diversity and global patterns remain poorly described. Here, we analyzed 285 ARGs in soils from 1012 sites across all continents, and created the first global atlas with the distributions of topsoil ARGs.

**Results:** We show that ARGs peaked in high latitude cold and boreal forests. Climatic seasonality and mobile genetic elements, associated with the transmission of antibiotic resistance, were also key drivers of their global distribution. Dominant ARGs were mainly related to multidrug resistance genes and efflux pump machineries. We further pinpointed the global hotspots of the diversity and proportions of soil ARGs.

**Conclusions:** Together, our work provides the foundation for a better understanding of the ecology and global distribution of the environmental soil antibiotic resistome.

## Background

Antibiotic resistance through the acquisition of antibiotic resistance genes (ARGs) evolves naturally via natural selection, and is a strategy used by bacteria to withstand the harmful effects of antibiotics released by bacteria and other organisms, playing a critical role in regulating microbial populations [1, 2]. The prevalence of ARGs in the environment has been triggered by the development and widespread use of antibiotics in human health care and animal production [3-6]. Soils are one of the most important reservoirs of ARGs on Earth (i.e., the soil antibiotic resistome) [1], and constitute a major pathway for the exchange of ARGs among bacteria, including major clinical pathogens [7, 8]. The accumulation of ARGs in soils has emerged as a public health concern due to their high sequence similarity to ARGs in human pathogens [7] and the potential of soil ARGs to reduce the effectiveness of antibiotics [9-10]. Moreover, the importance and implications of the accumulation of ARGs in soils have fostered multiple investigations aiming to understand the environmental factors controlling their abundance and diversity from local to global scales [11, 12]. In recent years, information on soil ARGs has increasingly become available at reference databases such as RefSoil+ [13]. However, there are still major unknowns associated with the global distributions of soil ARGs, and with the environmental drivers of their abundance and diversity. For example, despite the recognized importance and risks of increased antimicrobial resistance under global change [5], we still lack global atlas with the distributions of soil ARG diversity and abundance. This information is lacking even for the most dominant individual ARGs found across global soils. This knowledge is essential to identify the global soil ARGs hotspots, to better understand the ecology and biogeography of ARG-associated soil microbial communities, and to guide management actions aimed at reducing antibiotic resistance associated infections.

Improving our understanding of soil-borne ARGs is fundamental for two main reasons. First, soil ARGs constitute important defense tools used by soil microbes to out-compete other microorganisms for essential soil resources (e.g., nutrients) [11]. Thus, learning more about soil-borne ARGs, which are essential components of the soil microbiome with important functional implications (e.g., microbial warfare) [11], is fundamental to better understand the ecology and global distribution of microbial traits worldwide, about which we currently know very little. Second, under certain circumstances (e.g., soils with high accumulation of ARGs and MGEs under strong selection pressure) [14], soil-borne ARGs could potentially be transmitted to important plant, human and animal pathogens [14], reducing our capacity to fight important diseases. Knowing the location of soil-borne ARGs hotspots could be of great help to anticipate these situations, and thus to prevent future potential threats to human, animal and plant health.

To address existing knowledge gaps related to topsoil ARGs, we conducted the largest and most comprehensive standardized global survey carried out to date (1012 locations across 35 countries from all continents; Fig. 1), and used a high throughput quantitative PCR approach [15] to characterize the richness (number of ARG phylotypes) and proportion of 285 individual topsoil (10 cm depth) ARGs encoding resistance to major categories of clinically relevant antibiotics (Supplementary Table 1). The locations surveyed include a wide range of terrestrial ecosystem types (croplands and natural ecosystems such as forests, grasslands and shrublands) and climatic regions (arid, temperate, tropical, continental, and polar; Supplementary Table 2), and capture a representative fraction of global environmental conditions (Supplementary Fig. 1). The standardized proportion of topsoil ARGs was determined as the average standardized (between 0 and 1) relative abundance (normalized by accounting for bacterial 16S rRNA gene) of 285 individual ARGs (see Methods). Using this approach, each ARG or MGE contributed equally to the final relative abundance (proportion) of ARGs or MGEs. The proportion of ARGs and MGEs calculated using this approach were highly correlated with the same variables calculated as the sum of all standardized (0-1) ARGs or MGEs (ρ =1.00; P < 0.001) or as the sum of the non-standardized relative abundance of all ARGs (ρ =0.89; P < 0.001) or MGEs (ρ =0.95; P < 0.0001).

**Figure 1.**
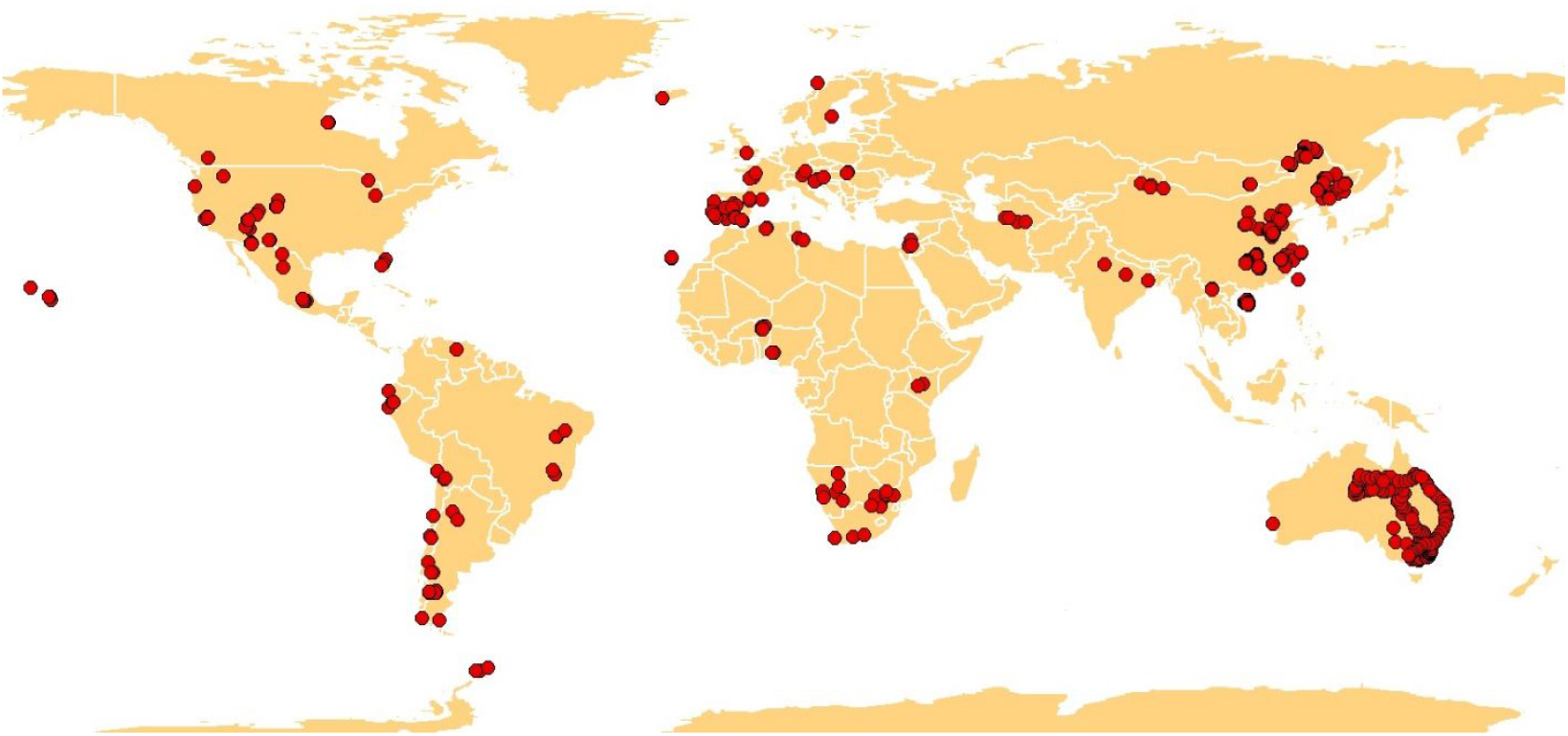
Location of the 1012 sites included in this study.

Using this unique survey, we: (i) identified the most dominant ARG types and mechanisms of action (e.g., protection, deactivation, efflux) in soils worldwide, (ii) investigated the main environmental factors (climate, soil properties, vegetation and proportion and richness of mobile genetic elements; MGEs) controlling the distribution of the proportion and diversity (richness) of topsoil ARGs, and (iii) generated an atlas identifying their global distribution across terrestrial ecosystems. We hypothesized that: (1) similar to what it has been reported for bacterial individual taxa [16], a few ARG types dominates soils across the globe; and (2) the richness and proportion of MGEs play an overriding role in controlling the global distribution of ARGs. We know that soils play an important role in the transmission of ARGs from one microbe to another through horizontal gene transfer [10-12], yet the relative importance of mobile genetic elements (MGEs) compared with other environmental factors in driving the global distributions of ARGs remains poorly understood. We also expect our global mapping effort to provide a new perspective of the potential global hotspots for the richness and proportion of ARGs.

## Materials and methods

### Global field survey and soil sample collection

Composite soil samples (from multiple soil cores) were collected from 1012 sites from 35 countries across all continents (Fig. 1). Surveyed plots were ∼0.1-0.25ha, and aim to provide information at the ecosystem level. These sites include a wide range of vegetation types (forests, grasslands and shrublands in natural ecosystems, and croplands; Supplementary Table 2) and climatic regions (arid, temperate, tropical, continental, and polar ecosystems). Croplands included rice, maize, soybean, tea and peanut, were mostly based in Asia and mainly fertilized using inorganic fertilizers. Also, to the best of our knowledge, these croplands have not been watered with reclaimed water. Mean annual precipitation and temperature in these locations ranged from 26 to 2347 mm and from -6.7 to 29.2ºC, respectively. Soil sample collection took place between 2012 and 2019. At each site, a topsoil sample (top ∼10 cm depth) was collected under the most common vegetation. At each location, we collected a composite soil sample based on multiple soil cores (10-15 cores) to account for the spatial heterogeneity within the surveyed plot. After field collection, each soil sample was separated into two sub-samples. One sub-sample was frozen at -20 ºC for molecular analyses while the other sub-sample was air-dried for chemical analyses. Soil pH and carbon ranged between 2.99-9.54 and 0.03-48.28%, respectively.

## ARGs and MGEs characterization

Soil DNA was extracted using the Powersoil® DNA Isolation Kit (MoBio Laboratories, Carlsbad, CA, USA) according to the manufacturer’s instructions. DNA was shipped to the University of Melbourne, where all samples were processed using the same standardized protocols. The relative abundance of 10 unique MGEs and 285 unique ARGs encoding resistance to all the major categories of antibiotics was obtained from all soil samples using the Waferegen SmartChip Real-Time PCR system (Fremont, CA, USA) [15]. This high throughput quantitative PCR technology is both a powerful tool to detect a wide spectrum of primer-specific ARGs and MGEs [12, 17, 18] and suitable to conduct comparative studies of antibiotic resistance [17]. It has also been widely used to investigate the relative abundance and diversity of ARGs in various environmental settings, including natural environments with limited human disturbance [12, 17-19]. Information on the primer sets used, and on the type and antibiotic resistance mechanism behind every ARG quantified is available in Supplementary Table 1.

We followed the PCR protocol described in [12]. In brief, the 100 nl reactions contained SensiMix SYBR No-ROX reagent (Bioline, London, UK), primers, DNA, and sterilized water. We included three analytical replicates for each soil sample and run. We used 5184-nanowell Smartchips (Wafergen, Fremont, CA, USA) including 296 primer sets, a calibrator (as 16S rRNA gene for the same DNA sample for all the chips) and a negative control. All primer sets used in this study have been validated to reduce the rate of false positives. The newest version of this high-throughput PCR method for ARG detection included 384 primer sets, the results obtained from the updates primer sets correlate well with old primers, suggesting that comparisons can still be made for samples analyzed using the old and new arrays [18]. To ensure reproducibility of our results, we used a high-precision, nanoliter-volume liquid handler (i.e. Smartchip MultiSample NanoDispenser) to process our samples. Amplification conditions were 95 °C for 10 min, followed by 40 cycles of 95 °C for 30 s and 60 °C for 30 s. The results were filtered following the next criteria: (i) a threshold cycle value (C_T_) of 31 was used as the detection limit; (ii) samples with more than two analytical replicates with a C_T_ less than 31 were regarded as positive quantification; and (iii) amplicons with multiple melting curve peaks were removed from the analysis.

We used the 2^−ΔCT^ method [20], where ΔC_T_ = (C_T detected ARGs_ − C_T 16S rRNA gene_), to calculate the relative abundances of ARGs and MGEs compared to the abundance of 16S rRNA gene in each soil sample. We then standardized the relative abundance of all individual ARGs and MGEs between 0 and 1. The proportion of ARGs and MGEs used in the main analyses from this paper was calculated as the average of the standardized relative abundance of all individual ARGs or MGEs.

## Environmental information

The coordinates and ecosystem type (grassland, shrubland, forest or cropland) of each location surveyed were recorded *in situ*. Information on the total annual temperature and precipitation (BIO1 and 12), as well as on their variability (BIO4 and 15) was obtained from the WorldClim v2 database at 1 km resolution [21] (https://www.worldclim.org/data/bioclim.html). The climatic variables included here are calculated as explained in [22], and are based on highly standardized, well-accepted and long-term used climatic variables[21, 22]. Soil pH was measured with a pH meter, in a 1:2.5 mass: volume soil and water suspension. Soil fine texture (% of fine fractions: clay + silt) and the concentration of soil total organic C and total N were measured using standardized methods[22, 23]. Soil C:N was calculated using the above information. Total organic C and N were highly correlated with each other (r = 0.90; P < 0.001), and therefore, only total organic C (but not N) was used in subsequent analyses.

## Statistical analyses

We used non-parametric PERMANOVA to test for significant differences (P < 0.05) in the richness [24], the proportion of ARGs and their community composition across biomes and continents (see Supplementary Table 2 for further details on our biome classification). PERMANOVA analyses were carried out using PRIMER v 6113 and PERMANOVA+ (PRIMER-E, Plymouth, UK) considering every composite sample/site as a replicate. Having more than one sample within each site would have been considered pseudo-replication as our question was related to compare the proportion and richness of soil ARGs across different ecosystem types globally (e.g., boreal vs. tropical forests) rather than to compare the proportion and richness of soil ARGs across sites within a given ecosystem type (e.g., two temperate forests). Environmental gradient designs [25] such as that used here are considered a powerful tool for detecting patterns in ecological responses, and generally outperform local replicated designs in terms of prediction success of responses at a large spatial scale [25]. PERMANOVA analyses are often apply to any situation where one wishes to analyze either univariate or multivariate data in response to either simple or complex experimental designs or models. The methods are particularly suited for non-parametric data.

We then used Structural Equation Modelling (SEM) [26] to identify the direct and indirect effects of space, climate, vegetation, MGEs and soil properties as drivers of the richness and proportion of soil ARGs (the main structure of our *a priori* model can be found in Supplementary Fig. 2; also see Supplementary Table 3 for further details on the predictors used and Supplementary Tables 4-5 for all direct associations considered). We grouped the different categories of predictors (climate, soil properties, vegetation and MGEs) in the same box for graphical simplicity. But please note that these boxes do not represent latent variables (Supplementary Fig. 2). Variables within these boxes are allowed to covary, with the exception of elevation and spatial dissimilarity, which constituted our degree of freedom (Supplementary Fig. 2). The most globally distributed vegetation types in our database (forests, shrublands, and grasslands) were included in our SEM as categorical variables with two levels: 1 (a given vegetation type) and 0 (remaining vegetation types). Since some of the variables introduced were not normally distributed, the probability that a path coefficient differs from zero was tested using bootstrap tests [27]. Bootstrapping is preferred to the classical maximum-likelihood estimation in these cases, because in bootstrapping, probability assessments are not based on an assumption that the data match a particular theoretical distribution. We then tested the goodness of fit of our model using the approach explained in refs. (26) and (27), which include information on the Chi-square test (χ^2^; the model has a good fit when 0 ≤ χ^2^ ≤ 2 and 0.05 < p ≤ 1.00), root mean square error of approximation (RMSEA; the model has a good fit when RMSEA 0 ≤ RMSEA ≤ 0.05 and 0.10 < p ≤ 1.00) and the Bollen-Stine bootstrap test (the model has a good fit when 0.10 < bootstrap p ≤ 1.00). Our model showed a solid goodness-of-fit, and therefore, a satisfactory fit to our data. SEM models were conducted with the software AMOS 20 (IBM SPSS Inc, Chicago, IL, USA). Finally, we also repeated our models using the subset of natural ecosystems (n = 802), which included the largest proportion of our data.

## Global mapping

Mapping of the proportion and richness of soil ARGs was done using a Random Forest algorithm [28]. Model fitting and prediction were done using ArcGIS Pro (ESRI, Redlands, California, U.S.A). Random forest algorithms create hundreds of ensembles of decision trees to create a model that can then be used for prediction [28, 29]. For each decision tree, a random set of points generated from the training data (here two thirds of the dataset) is used to calculate a prediction that then contributes to the overall outcome. To ensure compatibility across datasets and variables, we calculated all maps at a spatial resolution of 0.25 degrees. We considered as predictors for: climate (mean annual temperature; mean annual precipitation; temperature seasonality; precipitation seasonality), land-use type (forests; grasslands; cropland; shrublands), vegetation cover, elevation, and soil variables (soil fine texture; carbon content; pH and C:N ratio). When generating our global maps, information on soil properties was collected from SoilGRIDS (https://soilgrids.org) and resampled to 0.25 degrees by using the mean value of all pixels contained in each cell. The same applied to elevation [21] and vegetation cover [30]. While our global survey cover most of the variability in environmental conditions found on Earth, we left out of our predictions locations of the planet with high uncertainty in our database (areas in grey in Supplementary Fig. 1). Our models returned an R^2^ of 0.92 (for richness) and 0.86 (for relative abundance). Further cross-validation of model fit was done with a randomized set of 10% of the samples. The resulting correlations between predicted and observed richness and proportion of soil ARGs were highly significant in both cases (P < 0.0001).

## Results

First, we investigated the proportion and richness of ARGs in soils worldwide. Using a histogram, we could infer that while most surveyed soils showed relatively low proportions of ARG, only a few soils had relatively high proportions of topsoil ARGs (Fig. 2A). Intermediate levels of ARG richness (40-60 out of 285 ARGs) were common across the soils surveyed (Fig. 2A). We also found that multidrug resistance genes, and efflux pump machineries, were the most dominant types of ARGs in soils globally (Fig. 3A). We further identified, for the first time, dominant ARGs that were abundant (within the top 20% most abundant ARGs) and ubiquitous (occurred at >50% of sites and were present in >2/3 of the global biomes in Supplementary Table 2) across global soils. We found 14 dominant ARGs in soils across the globe (Fig. 2B). The beta-lactamase gene *fox5* and the multidrug resistance genes *oprJ, oprD*, and *acrA-05* were the most dominant ARGs in global soils. These dominant ARGs are less likely artifacts as only ARG assays with perfect single peak in melting curves and high amplification efficiency are retained in our results. We also included a water sample in each Smartchip as a negative control, and each DNA sample was run with three technical replicates to reduce any false positive/negative detections.

**Figure 2.**
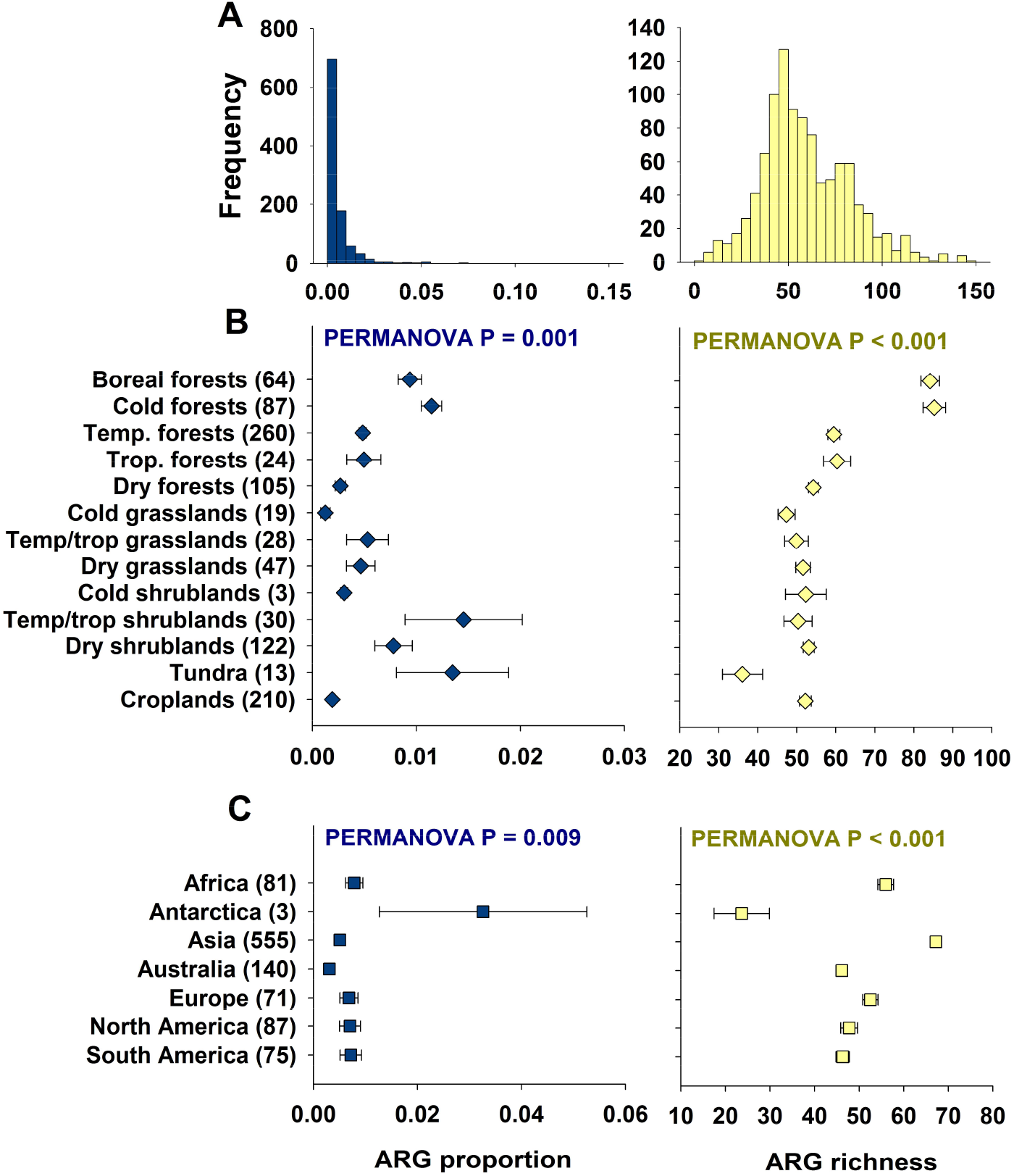
Proportion and richness of antibiotic resistance genes (ARGs) across global biomes and continents. Panel (A) includes the distribution histogram for the proportion and richness of ARGs. Panels (B) and (C) include the mean (± SE, number of sites/ecosystems in brackets) of the proportion and richness of soil ARGs across global biomes and continents, respectively. Each site is considered a statistical replicate in the PERMANOVA analyses. The proportion of ARGs was determined as the average standardized relative abundance of 285 individual ARGs. Details of the global biome classification can be found in Supplementary Table 2. Temp. = Temperate. Trop. = Tropical. We grouped our data to ensure high enough resolution for all ecosystem sub-types.

**Figure 3.**
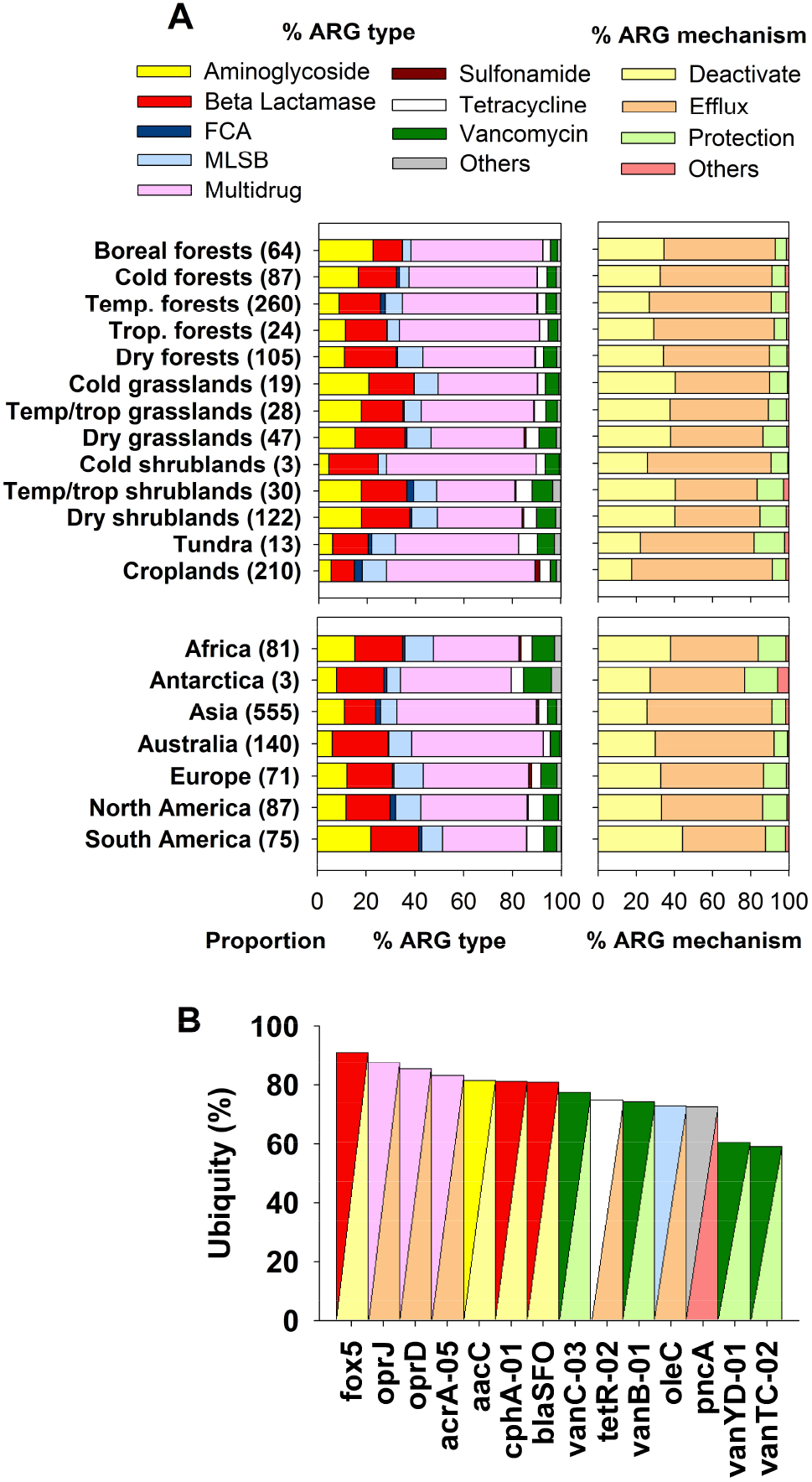
ARG composition across global biomes and continents. Panel (A) shows the proportion (%) of ARG types across global biomes and continents (*n* for each biome in brackets). Panel (B) shows the ubiquity (%) of dominant ARGs. Dominant ARGs are those that are abundant (top 20% of abundance), ubiquitous (occurred at >50% of sites) and present in at least 2/3 of the biomes surveyed. Details on the global biome classification used can be found in Supplementary Table 2. Temp. = Temperate and Trop. = Tropical. An additional visualization of the community composition of all ARGs across global biomes and continents can be found in Fig. 4.

Ecosystem biomes had a large influence on the proportion and richness of ARGs (Fig. 4). First, we showed that tundra ecosystems, boreal forests, other cold forests, and shrublands were positively associated with the proportion of soil ARGs globally (Fig. 2B; see Supplementary Table 2 for a biome classification). Boreal, cold, temperate and tropical forests were positively associated with the richness of ARGs in soils (Fig. 1B). We did not find significant differences in the richness of ARGs between croplands and other biomes (Fig. 2B).

**Figure 4.**
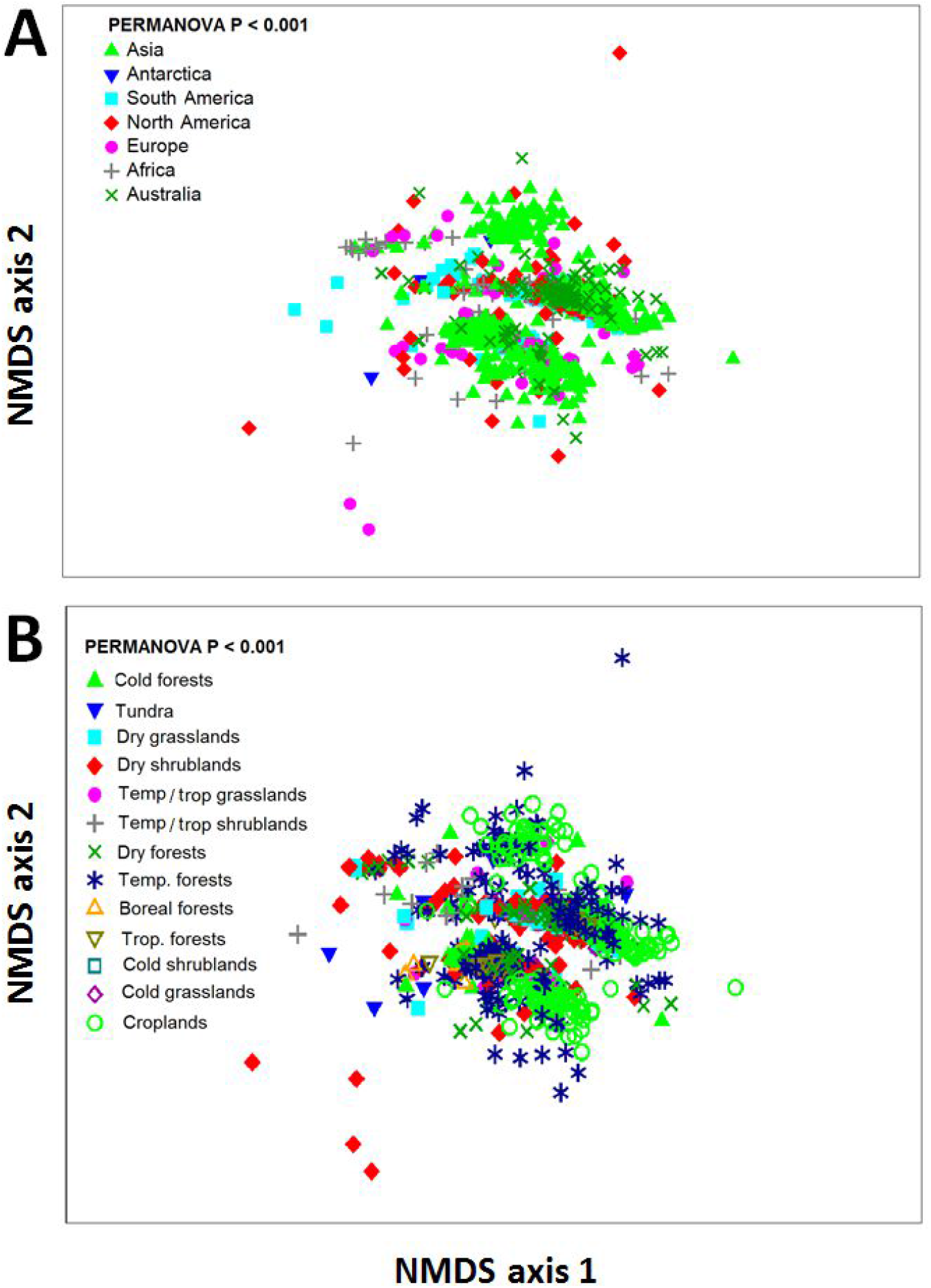
Community composition of ARGs across continents (A) and global biomes (B). NMDS analysis (Bray-Curtis) summarizing the community composition information (relative abundance) of 285 ARGs across different continents and global biomes (see Supplementary Table 2).

We then aimed to identify the major environmental factors associated with the distribution of the proportion and richness of soil ARGs. Our SEM revealed that the proportion of soil MGEs was the most important factor positively associated with the proportion of soil ARGs (Figs. 5A and C; Bootstrap *P* = 0.001; Supplementary Tables 3-4); this relationship was more important than that found for key environmental factors such as location, climate, vegetation, and soil properties (Fig. 5A; Supplementary Tables 3-4). Our SEM provided further evidence of direct and significant negative correlations between the proportion of soil ARGs with mean annual temperature and temperature seasonality, and of direct and significant positive relationship between the proportion of soil ARGs and precipitation seasonality (Fig. 5A). Similarly, the richness of MGEs was highly positively correlated with that of ARGs (Fig. 5B). We also found important indirect associations of MGE richness with ARG richness via changes in pH and plant cover (Fig. 5B). Spearman correlations between environmental factors and ARGs are available in Fig. 6.

**Figure 5.**
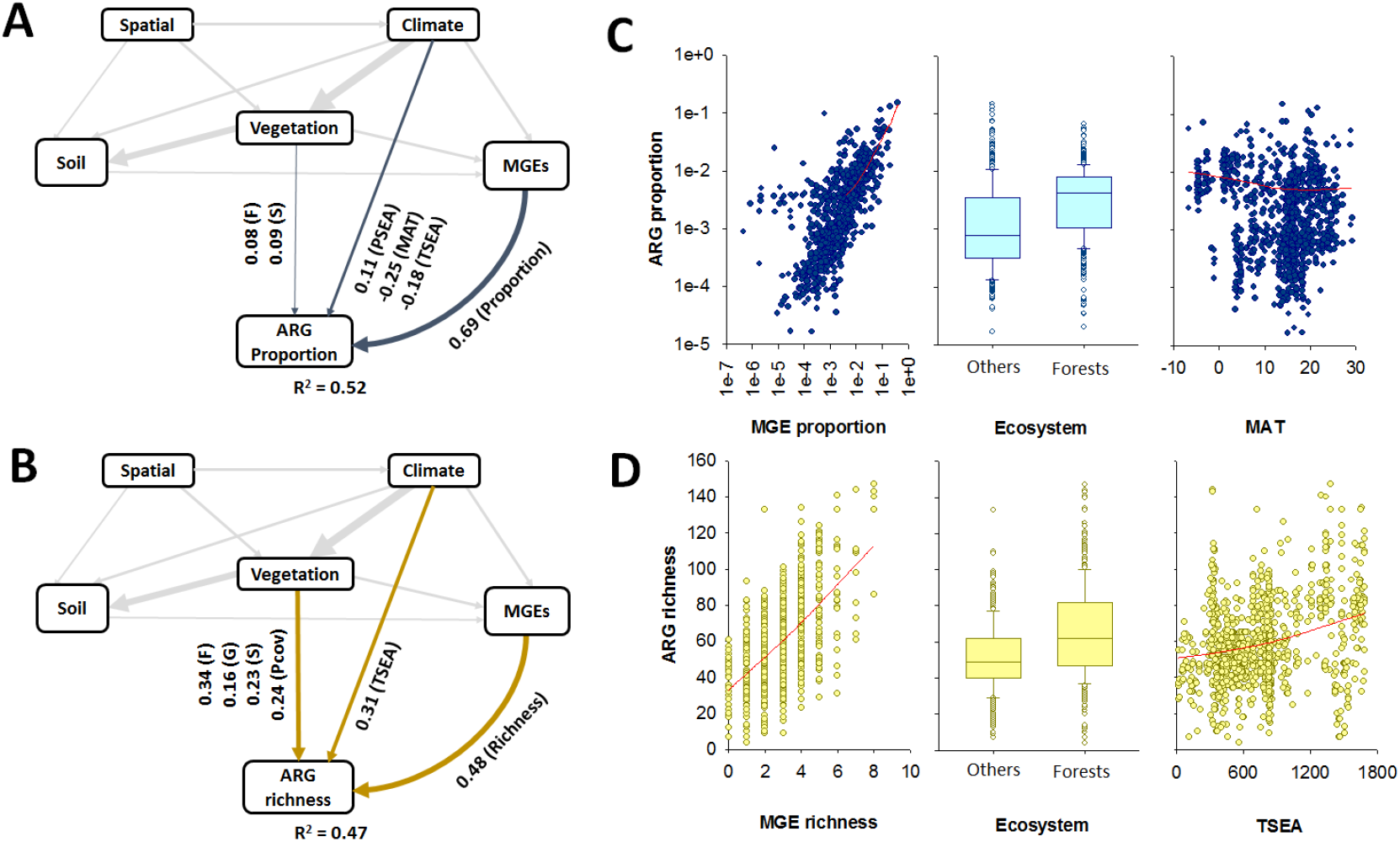
Drivers of the proportion (A) and richness (B) of topsoil ARGs globally. Panels A and B include structural equation models assessing the direct and indirect effects of environmental factors on the proportion and richness of ARGs. The proportion of ARGs was determined as the average standardized relative abundance of 285 individual ARGs. We grouped the different categories of predictors (climate, soil properties, vegetation and MGEs) in the same box for graphical simplicity (these boxes do not represent latent variables). Variables within these boxes are allowed to covary, with the exception of elevation and spatial dissimilarity, which constituted our degree of freedom. Numbers adjacent to arrows are indicative of the effect size of the relationship. Only significant effects (P < 0.05) are plotted. Supplementary Tables 4-5 shows the full SEM. F = Forests; G = Grasslands; S = Shrublands. MAT = mean annual temperature. PSEA = precipitation seasonality. TSEA = temperature seasonality. There was a non-significant deviation of the data from the model (χ2 =0.10, df = 1; *P* = 0.75; RMSEA *P* = 0.93; Bootstrap *P* = 0.71). Panel C includes selected scatter and box-plots showing the regression between environmental factors and soil ARGs. Red lines are Loess regressions. MGEs includes both richness and proportions. Y-axis in Panel C is shown in log-scale. Units and acronyms are available in Supplementary Table 3.

**Figure 6.**
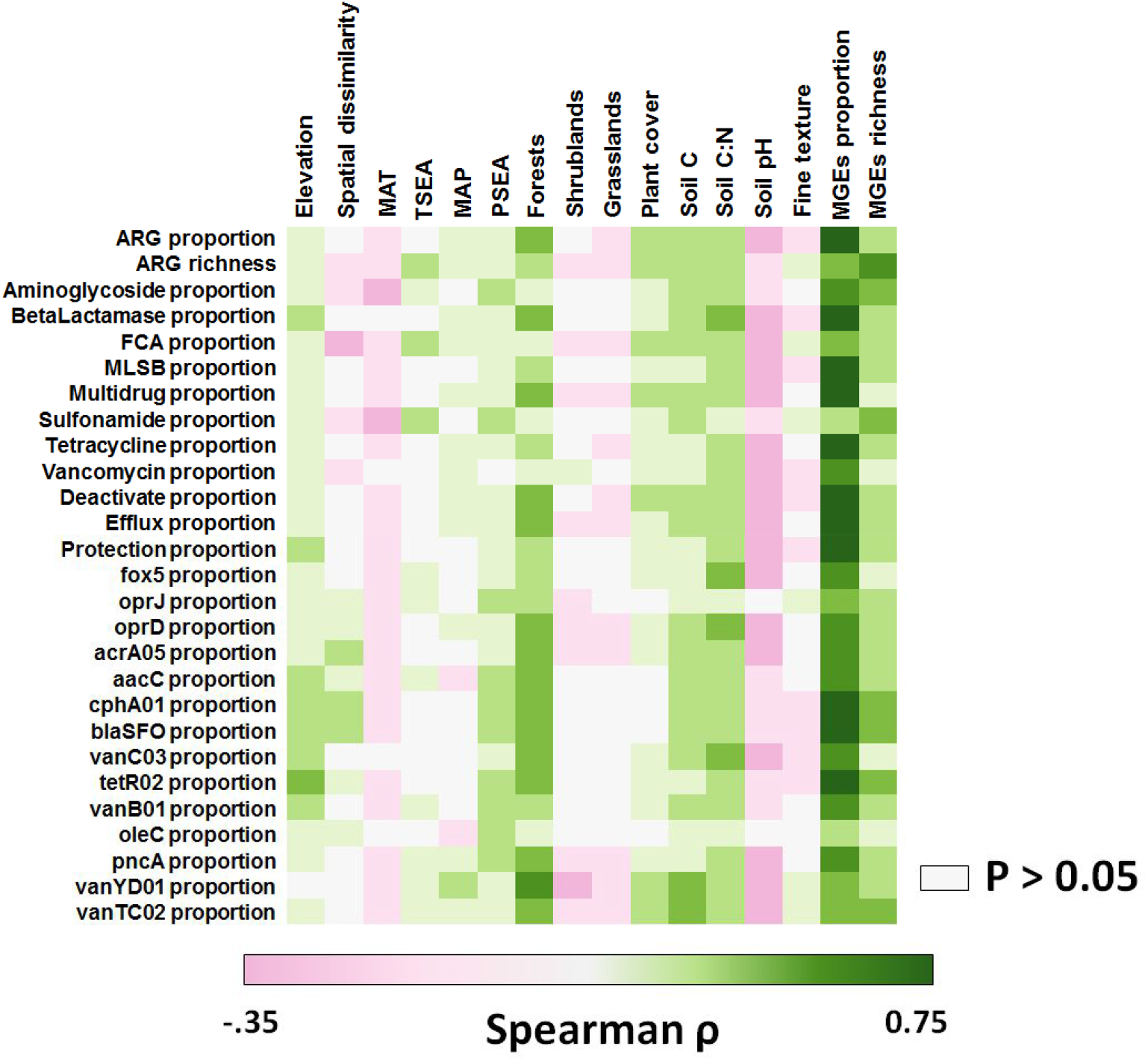
Spearman rank correlations between environmental predictors and the proportion and richness of soil ARGs. Acronyms are available in Supplementary Table 3. MAT = Mean annual temperature. MAP = Mean annual precipitation. PSEA = Precipitation seasonality. TSEA = Temperature seasonality. Non-significant correlation are plotted in gray.

We then created the first collection of maps for current distribution of the richness and proportions of soil ARGs (Fig. 7). Our atlas supported previous results in Fig. 1, and indicated that soils from cold/boreal forests in North America/Asia were associated with intermediate-high proportions of ARGs (Fig.7). High latitudinal regions of North America and Asia, as well as tropical and subtropical regions in South America, Africa and Asia were found to be the most important hotspots of the richness of soil ARGs (Fig. 7B).

**Figure 7.**
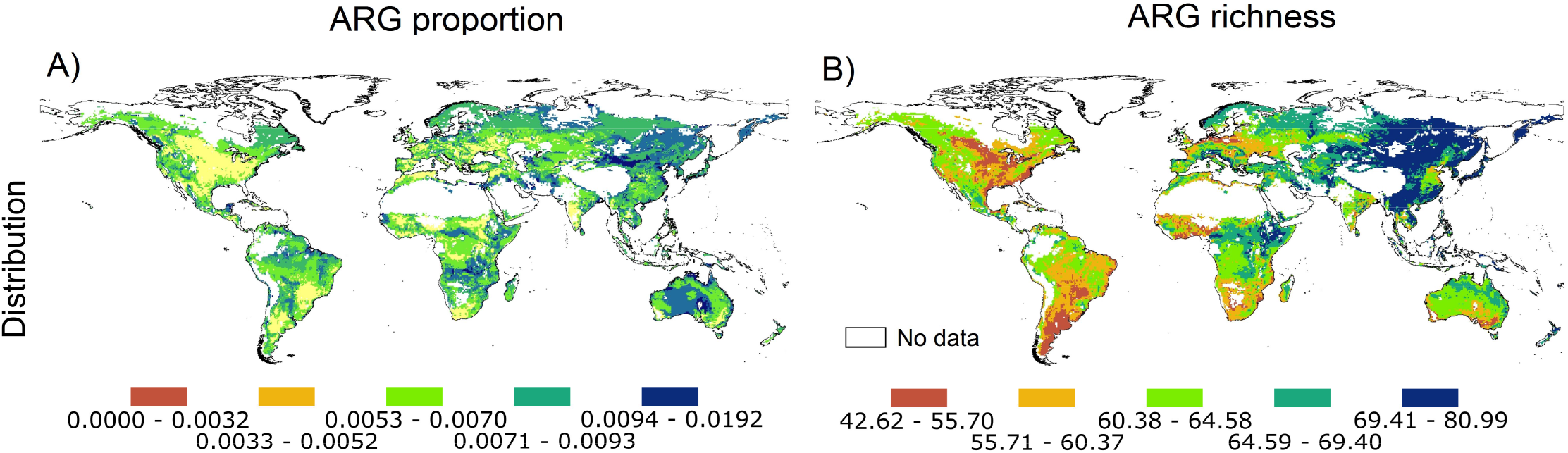
A global atlas of the distribution of topsoil ARGs. Panels (A) and (B) represent the present global distribution of the proportion and richness of soil ARGs, respectively. Numbers associated with the legend of this figure show standardized proportions and richness of topsoil ARGs. The proportion of ARGs was determined as the average standardized proportion of 285 individual ARGs. Our models returned an R^2^ of 0.92 (for richness) and 0.86 (for relative abundance). Outlier regions in our global survey (>97.5% quantile of the Chi-squared distribution) were not considered in our global atlas and are plotted in white as no data (see Supplementary Fig. 1).

## Discussion

Our study provides the first atlas showing the global distribution of environmental ARGs in soils, through conducting a global field survey of 1012 sites across 35 countries from all continents. We pinpointed the major global hotspots of soil ARGs and identified global drivers for diversity and abundance of the dominant soil ARGs. This study represents an important advancement in our understanding of the ecology, biogeography and potential changes in soil-borne ARGs in a changing world, which is integral to increase our capacity to address future health crises driven by antibiotic resistant infections.

Our study shows that, in general, soils support relatively low ARG proportions, with only a few soils supporting relatively high proportions of topsoil ARGs (Fig. 2A). On the contrary, according to our histogram, intermediate levels of ARG richness are common in soils worldwide (Fig. 2A). Multidrug resistance genes, and efflux pump machineries, are the most dominant ARG types and antibiotic resistance mechanisms of action found in soils across the globe (Figs. 3A and 4). These ARGs are especially effective against a broad spectrum of antibiotics because they allow bacteria to pump antibiotic peptides out of their cells [37]. Efflux pumps, like *oprJ* and *oprD* revealed in this study, are evolutionarily ancient and a very common antibiotic resistance mechanism in pristine ecosystems [37, 38]. Multidrug efflux pumps are commonly intrinsically encoded by chromosome and exhibit different functions with physiological and ecological significances that go beyond their activity as antibiotic resistance elements. In fact, multidrug efflux pumps have a wide range of substrate and their original function was not, in most cases, to resist to antibiotics [38]. Their physiological roles also involve regulating intracellular pH, transporting quorum sensing molecules and enhancing bacterial pathogenicity. Our results are in agreement with previous local and experimental work highlighting the dominance of this type of ARGs in soils [37, 39], and support the many studies that have demonstrated the ubiquity of ARGs in terrestrial ecosystems with contrasting level of anthropogenic disturbance, from pristine to croplands [40-42].

We further show that only 14 ARGs can be considered dominant and ubiquitous in soils worldwide, including the beta-lactamase gene *fox5* and the multidrug resistance genes *oprJ, oprD*, and *acrA-05*. These dominant ARGs were present in all biomes (Supplementary Table 2). The only exception was *oleC*, which was found in all biomes except in cold shrublands. The potential public health risks associated with these ubiquitous ARGs should be interpreted with caution, as these ubiquitous soil ARGs are commonly involved in basic process in bacterial physiology and should be regarded as a potential risk only if they are captured by transferable genetic elements [43]. However, when these genes are subject to a high antibiotic load (e.g., in farms or in natural ecosystems where antibiotic concentrations are locally high; 44), they are more likely to become relevant for resistance development. Indeed, the spread of faecal matter across the globe via animal waste, sewage effluents and birds transporting microbes from urban habitats, has contributed to the ARG dissemination in farmland soils and estuaries [45-47]. Even so, it should be noted that there are extremely stringent bottlenecks for the transfer of ARGs from soil bacterial hosts to human pathogens [48], especially in natural habitats that are rarely colonized by human pathogens. The abundant and ubiquitous ARGs identified in our study are clear targets to further investigate the potential global contribution of soils in increasing the resistance of microbial pathogens to several antibiotics. Taken together, these results provide global insights into the most common types of ARGs found in soils globally.

We then used Structural Equation Modeling (SEM; Supplementary Fig. 2) to generate a system-level understanding of the most important ecological factors controlling the distribution of topsoil ARGs across the globe (Fig. 5; Supplementary Fig. 2 and Supplementary Tables 3-5). Our results suggest that the proportion of soil MGEs (see Materials and Methods) is, by far, the most important factor associated (positively) with the proportion of soil ARGs (Fig. 5; Bootstrap *P* = 0.001; Supplementary Tables 3-4). Moreover, our SEM shows for the first time that the direct relationship between the proportion of soil-borne MGEs and that of ARGs is far more important than the effects of other essential environmental factors such as location, climate, vegetation, and soil properties (Fig. 5A; Supplementary Tables 3-4). For example, the direct relationship between the proportion of soil MGEs and that of ARGs is three times more important than that associated with mean annual temperature, and between 18 and 230 times more important than that linked to soil C:N ratio and soil C, respectively (Fig. 5A; Supplementary Tables 3-4). Similarly, the richness of soil MGEs was also the most important factor associated with the richness of ARGs (Fig. 5B), which is consistent with previous findings from a regional-scale study of Chinese forest ecosystems [12]. Here, we further show some novel indirect positive associations between ARG richness with both soil pH and plant cover via increasing richness of MGEs (Supplementary Table 5). Additional analyses showed that the proportion of soil MGEs had the strongest correlation with the relative abundance of multiple ARG types and mechanisms of action, and with the abundance of the most dominant individual ARGs (Fig. 6). MGEs (including plasmids, integrons and transposons) are common in soils [12], can transfer genetic information from one species or replicon to another, and allow ARGs to efficiently disperse across different organisms [14]. They can also potentially facilitate the transfer of important ARGs from soil microorganisms to clinically important human pathogens [49]. The importance of MGEs was maintained even when removing from our analyses the subset of locations belonging to croplands (Supplementary Fig. 3). Our findings are therefore important because they provide evidence of the potential capacity of soils to contribute to the rapid spread of genes associated with the resistance to medically relevant antibiotics via horizontal gene transfer mediated by MGEs. This knowledge further contributes to better understanding the rapidly increasing amount of information on soil ARGs (e.g., via RefSoil+) [13]. It is also important to note that the strong correlations of ARGs and MGEs implicate the genetic potential of ARGs transfer, but the frequency of horizontal gene transfer in soil is generally low, as revealed by the evidence that the composition of soil resistome is correlated with the taxonomic composition of bacteria [43].

Tundra ecosystems (nine locations in Antarctica, Chile, and Iceland), boreal forests (Northern Hemisphere high latitude forests), other cold forests, and shrublands had the greatest proportion of soil ARGs globally (Fig. 2B; see Supplementary Table 2 to a biome classification). Soils in these ecosystems have, on average, between two- and nine-times higher proportions of soil ARGs than soils from other ecosystems. Remarkably, Antarctica, which was represented by only three sites near each other, included the soils with the highest proportions of ARGs (Fig. 2C). These results agree with those from local studies observing an accumulation of ARGs in Arctic ecosystems [19, 50]. Our SEM provided further evidence of direct and significant negative associations between mean annual temperature and temperature seasonality with the proportion of soil ARGs. We also found a direct and significant positive relationship between the proportion of soil ARGs and precipitation seasonality, an environmental condition shared by cold deserts such as those from Antarctica and many temperate shrublands [51, 52]. These relationships were still found when we focused on samples from natural ecosystems, and removed those from croplands (Supplementary Fig. 3). Together, our results show that a greater proportion of soil ARGs are found in extreme environments and highlight potential co-evaluative mechanisms aiming to provide resistance genes and adapt to harsh environments. Microbial antibiotics and extreme cold temperatures are known to cause similar types of damage in cellular components [53]. Consequently, soil microbes might use ARGs to withstand both types of stressors, and this may explain the patterns of ARG proportion observed in cold ecosystems [11, 53]. Interestingly, although we also found a positive correlation between soil C:N ratio and the proportion of soil ARGs (Fig. 6), as previously reported by studies based on shotgun sequencing [11], this positive association vanished when we considered environmental factors such as location, climate, vegetation and soil properties in our SEM. The combination of these factors have not been previously considered as predictors of soil ARG abundance and diversity at a global scale. As previously reported [11], we also found a correlation between mean annual precipitation and the proportion of soil ARGs (Fig. 6). However, this association was only evident when croplands were removed from our analyses (Supplementary Fig. 3). Cropland ecosystems are often irrigated, and this could have masked the importance of precipitation for ARGs when all data were analyzed together.

Boreal, cold, temperate and tropical forests supported the highest richness of ARGs in soils (Fig. 2B). Remarkably, on average, boreal and cold forests supported a 64% higher ARG richness than all the other ecosystems (Fig. 2B). Our analyses further demonstrate the contribution of forest biomes to the diversity of topsoil ARGs globally (Figs. 5B and D, and 6). Croplands did not show significantly different levels of ARG richness compared with most other biomes. However, we would like to stress that most croplands in the present dataset are from Asia and that future studies would need to better address the global impact of agriculture on soil ARGs. Vegetation structure was also an important predictor of the community composition of individual ARGs (Figs. 6). Our findings indicate, therefore, that any land use promoting reforestation or deforestation [54] may have important consequences for the global management of soil ARGs. Our SEM analyses further identified a direct and significant positive association between soil ARG richness and temperature seasonality (Fig. 5B and D). These results were consistent when removing the subset of locations belonging to croplands (Supplementary Fig. 3). However, after removing croplands, we found that many vegetation impacts on ARG richness were driven via changes in soil pH (the major driver of soil bacterial diversity [11]), and that precipitation seasonality also positively influenced the richness of ARGs in exclusively non-cropland ecosystems. Finally, we found a positive and significant correlation between the fungal-to-bacterial ratio and the richness and proportion of soil ARGs at those sites where this comparison was possible (r > 0.299; P < 0.005; n = 87). Many fungal species are known to produce antibiotics [19], supporting the positive association with the richness of ARGs across soils reported here and elsewhere [11]. We also considered the possibility that human influence, as measured with the Global Human Influence index [55] (see Methods), could influence our results. However, we could not find any significant correlation between ARG richness and this index of human influence (*p* > 0.05; n = 1012).

We generated the first atlas for the current distribution of the richness and proportions of soil ARGs, a necessary step to identify topsoil ARG hotspots and to predict sources of potential resistance to antibiotics associated with soil ARG reservoirs. Our atlas suggests that soils from cold and boreal forests in North America and Asia support intermediate to high proportions of ARGs (Fig. 7A), with similar proportions being found in highly seasonal arid regions across the globe (Fig. 7A; Supplementary Fig. 4A). Our findings further suggest that soils from high latitudinal regions of North America and Asia, as well as tropical and subtropical regions in South America, Africa and Asia are the most important hotspots for the richness of soil ARGs (Fig. 7B). Many of these locations correspond to forest environments, and regions with high temperature seasonality (Fig. 7B; Supplementary Fig. 4B). Soils from Asia had some of the highest ARG richness, matching regions with the highest current and forecasted human casualties associated with antibiotic resistance, and with some of the largest rates of antibiotic applications for animal production on Earth [46]. Moreover, soils from highly populated areas in Australia showed relatively low proportion and richness of soil ARGs, matching areas with the lowest human casualties associated with antibiotic resistance [56]. These areas are also some of the most (e.g., China) and least (e.g., Australia) populated regions on Earth. Of course, soil ARGs are not necessarily directly implicated in these casualties. However, they potentially support a reservoir of multiple ARG types and defense mechanisms that can be acquired by human pathogens, increasing their virulence or incidence in areas already severely affected by antibiotic resistance.

## Conclusions

Our results significantly advance our knowledge of the ecology, biogeography and environmental factors associated with environmental soil-borne ARGs. We generated the first atlas of the global distribution of soil ARGs, pinpointing the major global hotspots of soil ARGs, and providing a comprehensive identification of global drivers for their diversity and proportions. We further provide a key database including standardized information on soil ARGs for >1000 globally distributed locations, which will be useful for soil microbial ecologists to test multiple hypotheses related to the ecology, biogeography and evolution of soil microbiomes. Together, our results pave the way for an improved understanding on the ecology and global distribution of key microbial traits such as those associated with the topsoil antibiotic resistome, and provide key insights to better manage the soil ARG pool and understand the future potential implications of soil-borne ARGs for microbial warfare and human health.

## Ethical Approval and Consent to participate

Not applicable.

## Consent for publication

All of the co-authors have read and approved the submitted version of the manuscript.

## Availability of data and materials

All the materials, raw data, and protocols used in the article are available upon request and without restriction. The data used in this article are available from Figshare (https://figshare.com/s/5640a4e375272e4eebf1).

## Authors’ contributions

M.D-B. developed the original idea of the analyses presented in the manuscript, with inputs from H-W.H. Fieldwork was done by all coauthors except N.E. and C.A.G. ARG data was produced, handled and curated by H-W.H., Y-G.Z., Q-L.C., S.D. and J-Z.H. Other lab analyses were done by M.D-B., F.T.M., B.G., V.O. and S.A. Global mapping was done by C.A.G. and N.E. Statistical modelling was done by M.D-B. The manuscript was written by M.D-B. with contributions from all co-authors.

## Acknowledgements

We thank the researchers involved in the CLIMIFUN, MUSGONET and BIODESERT projects for collection of field data and soil samples. We thank Drs. Cesar Plaza, Ana Rey and Asunción de los Ríos for their valuable comments on a previous draft of our manuscript. We also thank Melissa Martin for revising the English of the manuscript, and Dr. Felipe Bastida for his help with lab analyses.

## Supplementary Information

Supplementary Figures 1-4

Supplementary Tables 1-5

