## Supplementary Information for "The global distribution and environmental drivers of the soil antibiotic resistome"

**This PDF file includes:**
Supplementary Figures 1-4
Supplementary Tables 1-5

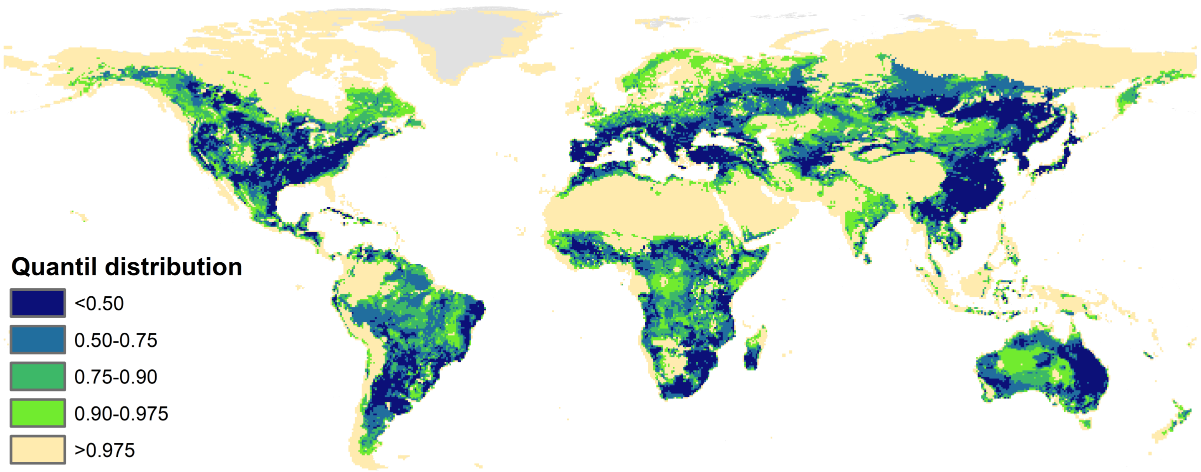


**Supplementary Figure 1. Extrapolation of uncertainties associated with the global survey used in this study.** Our map used the spatial accuracy indicator regarding the quantile distribution of the Chi-squared distribution with ten degrees of freedom based on the Mahalanobis multidimensional distance (corresponding to vegetation cover, annual precipitation, precipitation seasonality, mean annual temperature, temperature seasonality, soil texture, elevation, pH, soil carbon, and C:N ratio). This analysis provides information on those environmental conditions and locations of the planets which are underrepresented in our survey. Our dataset covers a wide range of the terrestrial environmental conditions found on Earth, being the rest considered as outliers (for this purpose an outlier is a location that has a value above the 97.5% quantile of the Chi-squared distribution).

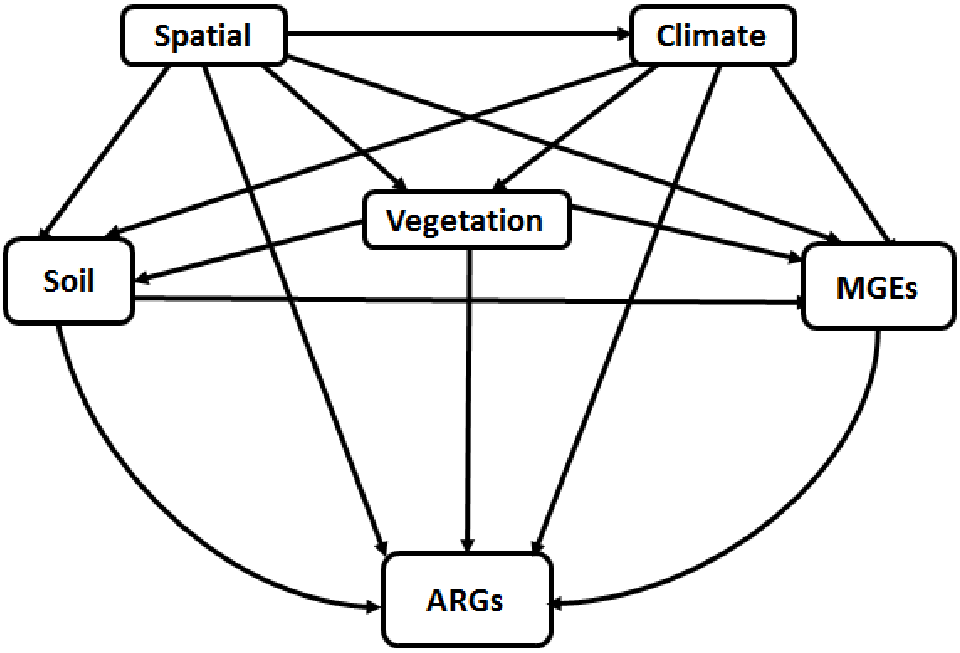


**Supplementary Figure 2. Main structure for the *a priori* structural equation model used in this study.** We grouped the different categories of predictors (space, climate, soil properties, vegetation and MGEs) in the same box for graphical simplicity (these boxes do not represent latent variables). See all considered direct associations in Supplementary Tables 4-5. Variables within these boxes are allowed to covary, with the exception of elevation and spatial dissimilarity, which constituted our degree of freedom.

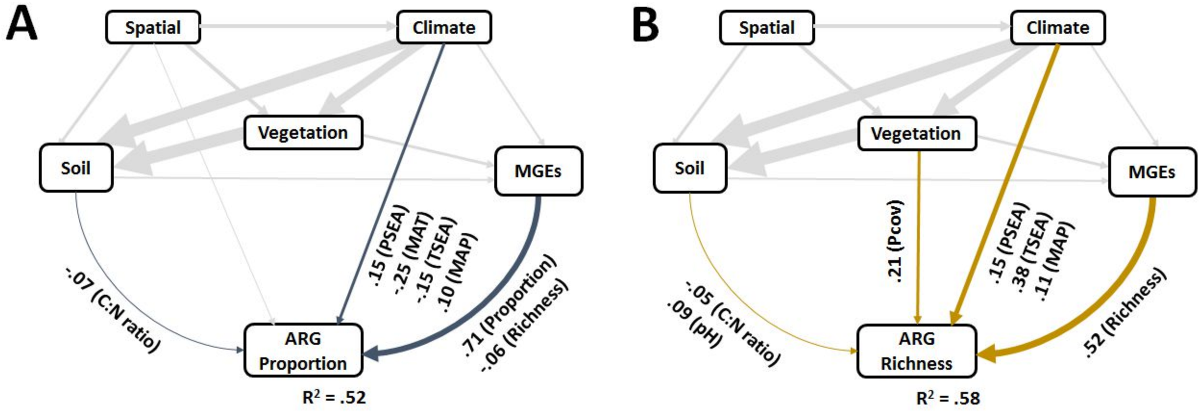


**Supplementary Figure 3.** Structural equation models assessing the direct and indirect effects of environmental factors on the proportion (A) and richness (B) of ARGs in natural ecosystems only (i.e. croplands excluded, n = 802). The proportion of ARGs was determined as the average standardized relative abundance of 285 individual ARGs. We grouped the different categories of predictors (climate, soil properties, vegetation and MGEs) in the same box for graphical simplicity (these boxes do not represent latent variables). Variables within these boxes are allowed to covary, with the exception of elevation and spatial dissimilarity, which constituted our degree of freedom. Numbers adjacent to arrows are indicative of the effect size of the relationship. Only significant paths ($P < 0.05$) are plotted. MGEs includes both richness and proportions.

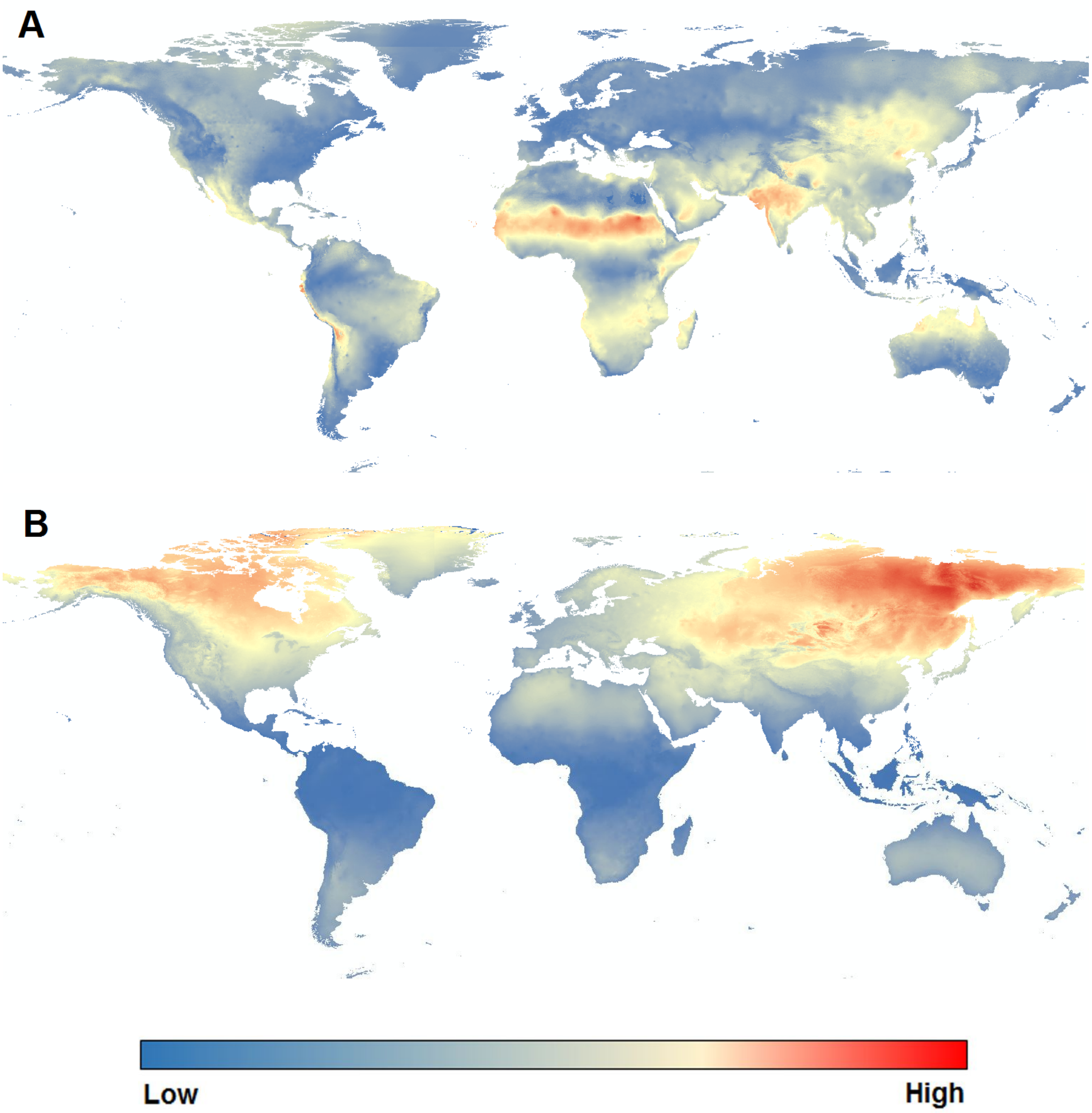


**Supplementary Figure 4.** Global precipitation (A) and temperature (B) seasonality maps used in our study (https://www.worldclim.org/data/index.html).

**Supplementary Table 1.** List of antibiotic resistance genes (ARGs) mobile genetic elements (MGEs) considered in this study.

| Gene Name | Type | Gene Classification | Resistance Mechanism | Forward Primer | Reverse Primer |
|---|---|---|---|---|---|
| 16S rRNA | 16S rRNA | 16S rRNA | Internal control | GGGTTGCGCTCGTTGC | ATGGYTGTCGTCAGCTCGTG |
| intI | MGES | MGE | integrase | GGCATCCAAGCAGCAAG | AAGCAGACTTGACCTGA |
| intI1 | MGES | MGE | integrase | CGAACGAGTGGCGGAGGGTG | TACCCGAGAGCTTGGCACCCA |
| tnpA-03 | MGES | MGE | transposase | AATTGATGCGGACGGCTTAA | TCACCAAACTGTTTATGGAGTCGTT |
| IS613 | MGES | MGE | transposase | AGGTTCGGACTCAATGCAACA | TTCAGCACATACCGCCTTGAT |
| tnpA-01 | MGES | MGE | transposase | CATCATCGGACGGACAGAATT | GTCGGAGATGTGGGTGTAGAAAGT |
| tnpA-04 | MGES | MGE | transposase | CCGATCACGGAAAGCTCAAG | GGCTCGCATGACTTCGAATC |
| tnpA-07 | MGES | MGE | transposase | GAAACCGATGCTACAATATCCAATTT | CAGCACCGTTTGCAGTGTAAG |
| tnpA-05 | MGES | MGE | transposase | GCCGCACTGTCGATTTTTATC | GCGGGATCTGCCACTTCTT |
| Tp614 | MGES | MGE | transposase | GGAAATCAACGGCATCCAGTT | CATCCATGCGCTTTTGTCTCT |
| tnpA-02 | MGES | MGE | transposase | GGGCGGGTCGATTGAAA | GTGGGCGGGATCTGCTT |
| catB3 | ARG | FCA | deactivate | GCACTCGATGCCTTCCAAAA | AGAGCCGATCCAAACGTCAT |
| catB8 | ARG | FCA | deactivate | CACTCGACGCCTTCCAAAG | CCGAGCCTATCCAGACATCATT |
| cfr | ARG | FCA | deactivate | GCAAAATTCAGAGCAAGTTACGAA | AAAATGACTCCCAACCTGCTTTAT |
| floR | ARG | FCA | efflux | ATTGTCTTCACGGTGTCCGTTA | CCGCGATGTCGTCGAACT |
| cmlA1-01 | ARG | FCA | efflux | TAGGAAGCATCGGAACGTTGAT | CAGACCGAGCACGACTGTTG |
| cmlA1-02 | ARG | FCA | efflux | AGGAAGCATCGGAACGTTGA | ACAGACCGAGCACGACTGTTG |
| cmx(A) | ARG | FCA | efflux | GCGATCGCCATCCTCTGT | TCGACACGGAGCCTTGGT |
| catA1 | ARG | FCA | deactivate | GGGTGAGTTTCACCAGTTTTGATT | CACCTTGTCGCCTTGCGTATA |
| qnrA | ARG | FCA | unknown | AGGATTTCTCACGCCAGGATT | CCGCTTTCAATGAAACTGCAA |
| aac | ARG | Aminoglycoside | deactivate | CCCTGCGTTGTGGCTATGT | TTGGCCACGCCAATCC |
| aacC1 | ARG | Aminoglycoside | deactivate | GGTCGTGAGTTCGGAGACGTA | GCAAGTTCCCGAGGTAATCG |
| aacC2 | ARG | Aminoglycoside | deactivate | ACGGCATTCTCGATTGCTTT | CCGAGCTTCACGTAAGCATTT |
| aacC4 | ARG | Aminoglycoside | deactivate | CGGCGTGGGACACGAT | AGGGAACCTTTGCCATCAACT |
| aac(6')I1 | ARG | Aminoglycoside | deactivate | GACCGGATTAAGGCCGATG | CTTGCCTTGATATTCAGTTTTTATAACCA |
| aacA/aphD | ARG | Aminoglycoside | deactivate | AGAGCCTTGGGAAGATGAAGTTT | TTGATCCATACCATAGACTATCTCATCA |
| aac(6')-Iy | ARG | Aminoglycoside | deactivate | GCTTTGCGGATGCCTCAAT | GGAGAACAAAAATACCTTCAAGGAAA |
| aac(6')-II | ARG | Aminoglycoside | deactivate | CGACCCGACTCCGAACAA | GCACGAATCCTGCCTTCTCA |
| aacC | ARG | Aminoglycoside | deactivate | CGTCACTTATTCGATGCCCTTAC | GTCGGGCGCGGCATA |
| aac(6')-Ib-01 | ARG | Aminoglycoside | deactivate | GTTTGAGAGGCAAGGTACCGTAA | GAATGCCTGGCGTGTTTGA |
| aac(6')-Ib-02 | ARG | Aminoglycoside | deactivate | CGTCGCCGAGCAACTTG | CGGTACCTTGCCTCTCAAACC |
| aadA5-01 | ARG | Aminoglycoside | deactivate | ATCACGATCTTGCGATTTTGCT | CTGCGGATGGGCCTAGAAG |
| aadA5-02 | ARG | Aminoglycoside | deactivate | GTTCTTGCTCTTGCTCGCATT | GATGCTCGGCAGGCAAAC |
| aac(6')-Ib-03 | ARG | Aminoglycoside | deactivate | AGAAGCACGCCCGACACTT | GCTCTCCATTCAGCATTGCA |

| | | | | | |
|---|---|---|---|---|---|
| aadA1 | ARG | Aminoglycoside | deactivate | AGCTAAGCGCGAACTGCAAT | TGGCTCGAAGATACCTGCAA |
| aadA2-01 | ARG | Aminoglycoside | deactivate | ACGGCTCCGCAGTGGAT | GGCCACAGTAACCAACAAATCA |
| aadA-01 | ARG | Aminoglycoside | deactivate | GTTGTGCACGACGACATCATT | GGCTCGAAGATACCTGCAAGAA |
| aadA2-02 | ARG | Aminoglycoside | deactivate | CTTGTCGTGCATGACGACATC | TCGAAGATACCCGCAAGAATG |
| aadD | ARG | Aminoglycoside | deactivate | CCGACAACATTTCTACCATCCTT | ACCGAAGCGCTCGTCGTATA |
| aadA2-03 | ARG | Aminoglycoside | deactivate | CAATGACATTCTTGCGGGTATC | GACCTACCAAGGCAACGCTATG |
| aadA9-01 | ARG | Aminoglycoside | deactivate | CGCGGCAAGCCTATCTTG | CAAATCAGCGACCGCAGACT |
| aadA-02 | ARG | Aminoglycoside | deactivate | CGAGATTCTCCGCGCTGTA | GCTGCCATTCTCCAAATTGC |
| aadA9-02 | ARG | Aminoglycoside | deactivate | GGATGCACGCTTGGATGAA | CCTCTAGCGGCCGGAGTATT |
| aadE | ARG | Aminoglycoside | deactivate | TACCTTATTGCCCTTGGAAGAGTTA | GGAACTATGTCCCTTTTAATTCTACAATCT |
| spcN-01 | ARG | Aminoglycoside | deactivate | AAAAGTTCGATGAAACACGCCTAT | TCCAGTGGTAGTCCCCGAATC |
| spcN-02 | ARG | Aminoglycoside | deactivate | CAGAATCTTCCTGAAAAGTTTGATGAA | CGCAGACACGCCGAATC |
| aadA-1-01 | ARG | Aminoglycoside | deactivate | AAAAGCCCGAAGAGGAACTTG | CATCTTTCACAAAGATGTTGCTGTCT |
| aph6ia | ARG | Aminoglycoside | deactivate | CCCATCCCATGTGTAAGGAAA | GCCACCGCTTCTGCTGTAC |
| aph(2')-Id-02 | ARG | Aminoglycoside | deactivate | TGAGCAGTATCATAAGTTGAGTGAAAAG | GACAGAACAATCAATCTCTATGGAATG |
| aph(2')-Id-01 | ARG | Aminoglycoside | deactivate | TAAGGATATACCGACAGTTTTGGAAA | TTTAATCCCTCTTCATACCAATCCATA |
| aph | ARG | Aminoglycoside | deactivate | TTTCAGCAAGTGGATCATGTTAAAAT | CCAAGCTGTTTCCACTGTTTTTC |
| aphA1 | ARG | Aminoglycoside | deactivate | TGAACAAGTCTGGAAAGAAATGCA | CCTATTAATTTCCCCTCGTCAAAAA |
| aadA-1-02 | ARG | Aminoglycoside | deactivate | CGGAATTGAAAAAACTGATCGAA | ATACCGGCTGTCCGTCATTT |
| str | ARG | Aminoglycoside | deactivate | AATGAGTTTTGGAGTGTCTCAACGTA | AATCAAAACCCCTATTAAAGCCAAT |
| strA | ARG | Aminoglycoside | deactivate | CCGGTGGCATTTGAGAAAAA | GTGGCTCAACCTGCGAAAAG |
| strB | ARG | Aminoglycoside | deactivate | GCTCGGTCGTGAGAACAATCT | CAATTTCGGTCGCCTGGTAGT |
| ampC-01 | ARG | Beta Lactamase | deactivate | TGGCGTATCGGGTCAATGT | CTCCACGGGCCAGTTGAG |
| ampC-02 | ARG | Beta Lactamase | deactivate | GCAGCACGCCCCGTAA | TGTACCCATGATGCGCGTACT |
| ampC-04 | ARG | Beta Lactamase | deactivate | TCCGGTGACGCGACAGA | CAGCACGCCGGTGAAAGT |
| ampC-05 | ARG | Beta Lactamase | deactivate | CTGTTCGAGCTGGGTTCTATAAGTAAA | CAGTATCTGGTCACCGGATCGT |
| ampC-06 | ARG | Beta Lactamase | deactivate | CCGCTCAAGCTGGACCATAC | CCATATCCTGCACGTTGGTTT |
| ampC-07 | ARG | Beta Lactamase | deactivate | CCGCCCAGAGCAAGGACTA | GCTCGACTTCACGCCGTAAG |
| ampC-09 | ARG | Beta Lactamase | deactivate | CAGCCGCTGATGAAAAAATATG | CAGCGAGCCCACTTCGA |
| ampC/blaDHA | ARG | Beta Lactamase | deactivate | TGGCCGCAGCAGAAAGA | CCGTTTTATGCACCCAGGAA |
| bla1 | ARG | Beta Lactamase | deactivate | GCAAGTTGAAGCGAAAGAAAAGA | TACCAGTATCAATCGCATATACACCTAA |
| bla-ACC-1 | ARG | Beta Lactamase | deactivate | CACACAGCTGATGGCTTATCTAAAA | AATAAACGCGATGGGTTCCA |
| blaCMY | ARG | Beta Lactamase | deactivate | CCGCGGCGAAATTAAGC | GCCACTGTTTGCCTGTCAGTT |
| blaCMY2-01 | ARG | Beta Lactamase | deactivate | AAAGCCTCAT GGGTGCATAAA | ATAGCTTTTGTTTGCCAGCATCA |
| blaCMY2-02 | ARG | Beta Lactamase | deactivate | GCGAGCAGCCTGAAGCA | CGGATGGGCTTGTCCTCTT |
| blaCTX-M-01 | ARG | Beta Lactamase | deactivate | GGAGGCGTGACGGCTTTT | TTCAGTGCGATCCAGACGAA |
| blaCTX-M-02 | ARG | Beta Lactamase | deactivate | GCCGCGGTGCTGAAGA | ATCGGATTATAGTTAACCAGGTCAGATTT |

| | | | | | |
|---|---|---|---|---|---|
| blaCTX-M-03 | ARG | Beta Lactamase | deactivate | CGATACCACCACGCCGTTA | GCATTGCCCAACGTCAGATT |
| blaCTX-M-04 | ARG | Beta Lactamase | deactivate | CTTGGCGTTGCGCTGAT | CGTTCATCGGCACGGTAGA |
| blaCTX-M-05 | ARG | Beta Lactamase | deactivate | GCGATAACGTGGCGATGAAT | GTCGAGACGGAACGTTTCGT |
| blaCTX-M-06 | ARG | Beta Lactamase | deactivate | CACAGTTGGTGACGTGGCTTAA | CTCCGCTGCCGGTTTTATC |
| blaGES | ARG | Beta Lactamase | deactivate | GCAATGTGCTCAACGTTCAAG | GTGCCTGAGTCAATTCTTTCAAAG |
| blaIMP-01 | ARG | Beta Lactamase | deactivate | AACACGGTTTGGTGGTTCTTGTA | GCGCTCCACAAACCAATTG |
| blaIMP-02 | ARG | Beta Lactamase | deactivate | AAGGCAGCATTTCCTCTCATTTT | GGATAGATCGAGAATTAAGCCACTCT |
| bla-L1 | ARG | Beta Lactamase | deactivate | CACCGGGTTACCAGCTGAAG | GCGAAGCTGCGCTTGTAGTC |
| blaMOX/blaCMY | ARG | Beta Lactamase | deactivate | CTATGTCAATGTGCCGAAGCA | GGCTTGTCCTCTTTCGAATAGC |
| blaOCH | ARG | Beta Lactamase | deactivate | GGCGACTTGCGCCGTAT | TTTTCTGCTCGGCCATGAG |
| blaOKP | ARG | Beta Lactamase | deactivate | GCCGCCATCACCATGAG | GGTGACGTTGTCACCGATCTG |
| blaOXA1/blaOXA30 | ARG | Beta Lactamase | deactivate | CGGATGGTTTGAAGGGTTTATTAT | TCTTGGCTTTTATGCTTGATGTTAA |
| blaOXA10-01 | ARG | Beta Lactamase | deactivate | CGCAATTATCGGCCTAGAAACT | TTGGCTTTCCGTCCCATTT |
| blaOXA10-02 | ARG | Beta Lactamase | deactivate | CGCAATTATCGGCCTAGAAACT | TTGGCTTTCCGTCCCATTT |
| blaOXY | ARG | Beta Lactamase | deactivate | CGTTCAGGCGGCAGGTT | GCCGCGATATAAGATTTGAGAATT |
| blaPAO | ARG | Beta Lactamase | deactivate | CGCCGTACAACCGGTGAT | GAAGTAATGCGGTTCTCCTTTCA |
| blaPER | ARG | Beta Lactamase | deactivate | TGCTGGTTGCTGTTTTTGTGA | CCTGCGCAATGATAGCTTCAT |
| blaPSE | ARG | Beta Lactamase | deactivate | TTGTGACCTATTCCCCTGTAATAGAA | TGCGAAGCACGCATCATC |
| blaROB | ARG | Beta Lactamase | deactivate | GCAAAGGCATGACGATTGC | CGCGCTGTTGTCGCTAAA |
| blaSFO | ARG | Beta Lactamase | deactivate | CCGCCGCCATCCAGTA | GGGCCGCCAAGATGCT |
| blaSHV-01 | ARG | Beta Lactamase | deactivate | TCCCATGATGAGCACCTTTAAA | TTCGTCACCGGCATCCA |
| blaSHV-02 | ARG | Beta Lactamase | deactivate | CTTTCCCATGATGAGCACCTTT | TCCTGCTGGCGATAGTGGAT |
| blaTEM | ARG | Beta Lactamase | deactivate | AGCATCTTACGGATGGCATGA | TCCTCCGATCGTTGTCAGAAGT |
| blaTLA | ARG | Beta Lactamase | deactivate | ACACTTTGCCATTGCTGTTTATGT | TGCAAATTTCGGCAATAATCTTT |
| blaVEB | ARG | Beta Lactamase | deactivate | CCCGATGCAAAGCGTTATG | GAAAGATTCCCTTTATCTATCTCAGACAA |
| blaVIM | ARG | Beta Lactamase | deactivate | GCACTTCTCGCGGAGATTG | CGACGGTGATGCGTACGTT |
| blaZ | ARG | Beta Lactamase | deactivate | GGAGATAAAGTAACAAATCCAGTTAGATATGA | TGCTTAATTTTCCATTTGCGATAAG |
| cepA | ARG | Beta Lactamase | deactivate | AGTTGCGCAGAACAGTCCTCTT | TCGTATCTTGCCCGTCGATAAT |
| cfiA | ARG | Beta Lactamase | deactivate | GCAGCGTTGCTGGACACA | GTTCGGGATAAACGTGGTGACT |
| cfxA | ARG | Beta Lactamase | deactivate | TCATTCCTCGTTCAAGTTTTCAGA | TGCAGCACCAAGAGGAGATGT |
| cphA-01 | ARG | Beta Lactamase | deactivate | GCGAGCTGCACAAGCTGAT | CGGCCCAGTCGCTCTTC |
| cphA-02 | ARG | Beta Lactamase | deactivate | GTGCTGATGGCGAGTTTCTG | GGTGTGGTAGTTGGTGTTGATCAC |
| fox5 | ARG | Beta Lactamase | deactivate | GGTTTGCCGCTGCAGTTC | GCGGCCAGGTGACCAA |
| mecA | ARG | Beta Lactamase | protection | GGTTACGGACAAGGTGAAATACTGAT | TGTCTTTTAATAAGTGAGGTGCGTTAATA |
| NDM1 | ARG | Beta Lactamase | deactivate | ATTAGCCGCTGCATTGAT | CATGTCGAGATAGGAAGTG |
| pbp | ARG | Beta Lactamase | protection | CCGGTGCCATTGGTTTAGA | AAAATAGCCGCCCCAAGATT |
| pbp2x | ARG | Beta Lactamase | protection | TTTCATAAGTATCTGGACATGGAAGAA | CCAAAGGAAACTTGCTTGAGATTAG |

| | | | | | |
|---|---|---|---|---|---|
| Pbp5 | ARG | Beta Lactamase | protection | GGCGAACTTCTAATTAATCCTATCCA | CGCCGATGACATTCTTCTTATCTT |
| penA | ARG | Beta Lactamase | protection | AGACGGTAACGTATAACTTTTTGAAAGA | GCGTGTAGCCGGCAATG |
| carB | ARG | MLSB | efflux | GGAGTGAGGCTGACCGTAGAAG | ATCGGCGAAACGCACAAA |
| ereA | ARG | MLSB | deactivate | CCTGTGGTACGGAGAATTCATGT | ACCGCATTCGCTTTGCTT |
| ereB | ARG | MLSB | deactivate | GCTTTATTTCAGGAGGCGGAAT | TTTTAAATGCCACAGCACAGAATC |
| erm(34) | ARG | MLSB | protection | GCGCGTTGACGACGATTT | TGGTCATACTCGACGGCTAGAAC |
| erm(35) | ARG | MLSB | protection | TTGAAAACGATGTTGCATTAAGTCA | TCTATAATCACAACTAACCACTTGAACGT |
| erm(36) | ARG | MLSB | protection | GGCGGACCGACTTGCAT | TCTGCGTTGACGACGGTTAC |
| ermA | ARG | MLSB | protection | TTGAGAAGGGATTTGCGAAAAG | ATATCCATCTCCACCATTAATAGTAAACC |
| ermA/ermTR | ARG | MLSB | protection | ACATTTTACCAAGGAACTTGTGGAA | GTGGCATGACATAAACCTTCATCA |
| ermB | ARG | MLSB | protection | TAAAGGGCATTTAACGACGAAACT | TTTATACCTCTGTTTGTTAGGGAATTGAA |
| ermC | ARG | MLSB | protection | TTTGAAATCGGCTCAGGAAAA | ATGGTCTATTTCAATGGCAGTTACG |
| ermF | ARG | MLSB | protection | CAGCTTTGGTTGAACATTTACGAA | AAATTCCTAAAATCACAACCGACAA |
| ermJ/ermD | ARG | MLSB | protection | GGACTCGGCAATGGTCAGAA | CCCCGAAACGCAATATAATGTT |
| ermK-01 | ARG | MLSB | protection | GTTTGATATTGGCATTGTCAGAGAAA | ACCATTGCCGAGTCCACTTT |
| ermK-02 | ARG | MLSB | protection | GAGCCGCAAGCCCCTTT | GTGTTTCATTTGACGCGGAGTAA |
| ermT-01 | ARG | MLSB | protection | GTTCACTAGCACTATTTTTAATGACAGAAGT | GAAGGGTGTCTTTTTAATACAATTAACGA |
| ermT-02 | ARG | MLSB | protection | GTAAAATCCCTAGAGAATACTTTCATCCA | TGAGTGATATTTTTGAAGGGTGTCTT |
| ermX | ARG | MLSB | protection | GCTCAGTGGTCCCCATGGT | ATCCCCCCGTCAACGTTT |
| ermY | ARG | MLSB | protection | TTGTCTTTGAAAGTGAAGCAACAGT | TAACGCTAGAGAACGATTTGTATTGAG |
| lmrA-01 | ARG | MLSB | efflux | TCGACGTGACCGTAGTGAACA | CGTGACTACCCAGGTGAGTTGA |
| lnuA-01 | ARG | MLSB | deactivate | TGACGCTCAACACACTCAAAAA | TTCATGCTTAAGTTCCATACGTGAA |
| lnuB-01 | ARG | MLSB | deactivate | TGAACATAATCCCCTCGTTTAAAGAT | TAATTGCCCTGTTTCATCGTAAATAA |
| lnuB-02 | ARG | MLSB | deactivate | AAAGGAGAAGGTGACCAATACTCTGA | GGAGCTACGTCAAACAACCAGTT |
| lnuC | ARG | MLSB | deactivate | TGGTCAATATAACAGATGTAAACCAGATTT | CACCCCAGCCACCATCAA |
| matA/mel | ARG | MLSB | efflux | TAGTAGGCAAGCTCGGTGTTGA | CCTGTGCTATTTTAAGCCTTGTTTCT |
| mdtA | ARG | MLSB | efflux | CCTAACGGGCGTGACTTCA | TTCACCTGTTTCAAGGGTCAAA |
| mefA | ARG | MLSB | efflux | CCGTAGCATTGGAACAGCTTTT | AAACGGAGTATAAGAGTGCTGCAA |
| mphA-01 | ARG | MLSB | deactivate | CTGACGCGCTCCGTGTT | GGTGGTGCATGGCGATCT |
| mphA-02 | ARG | MLSB | deactivate | TGATGACCCTGCCATCGA | TTCGCGAGCCCCTCTTC |
| mphB | ARG | MLSB | deactivate | CGCAGCGCTTGATCTTGTAG | TTACTGCATCCATACGCTGCTT |
| mphC | ARG | MLSB | deactivate | CGTTTGAAGTACCGAATTGGAAA | GCTGCGGGTTTGCCTGTA |
| msrA-01 | ARG | MLSB | efflux | CTGCTAACACAAGTACGATTCCAAAT | TCAAGTAAAGTTGTCTTACCTACACCATT |
| msrC-01 | ARG | MLSB | efflux | TCAGACCGGATCGGTTGTC | CCTATTTTTTGGAGTCTTCTCTCTAATGTT |
| oleC | ARG | MLSB | efflux | CCCGGAGTCGATGTTCGA | GCCGAAGACGTACACGAACAG |
| pikR1 | ARG | MLSB | protection | TCGACATGCGTGACGAGATT | CCGCGAATTAGGCCAGAA |
| pikR2 | ARG | MLSB | protection | TCGTGGGCCAGGTGAAGA | TTCCCCTTGCCGGTGAA |

| | | | | | |
|---|---|---|---|---|---|
| vatB-01 | ARG | MLSB | deactivate | GGAAAAAGCAACTCCATCTCTTGA | TCCTGGCATAACAGTAACATTCTGA |
| vatB-02 | ARG | MLSB | deactivate | TTGGGAAAAAGCAACTCCATCT | CAATCCACACATCATTTCCAACA |
| vatC-01 | ARG | MLSB | deactivate | CGGAAATTGGGAACGATGTT | GCAATAATAGCCCCGTTTCCTA |
| vatC-02 | ARG | MLSB | deactivate | CGATGTTTGGATTGGACGAGAT | GCTGCAATAATAGCCCCGTTT |
| vatE-01 | ARG | MLSB | deactivate | GGTGCCATTATCGGAGCAAAT | TTGGATTGCCACCGACAAT |
| vatE-02 | ARG | MLSB | deactivate | GACCGTCCTACCAGGCGTAA | TTGGATTGCCACCGACAATT |
| vgaA-01 | ARG | MLSB | efflux | CGAGTATTGTGGAAAGCAGCTAGTT | CCCGTACCGTTAGAGCCGATA |
| vgaA-02 | ARG | MLSB | efflux | GACGGGTATTGTGGAAAGCAA | TTTCCTGTACCATTAGATCCGATAATT |
| vgb-01 | ARG | MLSB | deactivate | AGGGAGGGTATCCATGCAGAT | ACCAAATGCGCCCGTTT |
| vgbB-01 | ARG | MLSB | efflux | CAGCCGGATTCTGGTCCTT | TACGATCTCCATTCAATTGGGTAAA |
| vgbB-02 | ARG | MLSB | deactivate | ATACGAGCTGCCTAATAAAGGATCTT | TGTGAACCACAGGGCATTATCA |
| sul2 | ARG | Sulfonamide | protection | TCATCTGCCAAACTCGTCGTTA | GTCAAAGAACGCCGCAATGT |
| sul1 | ARG | Sulfonamide | protection | CAGCGCTATGCGCTCAAG | ATCCCGCTGCGCTGAGT |
| sulA/folP-01 | ARG | Sulfonamide | protection | CAGGCTCGTAAATTGATAGCAGAAG | CTTTCCTTGCGAATCGCTTT |
| sulA/folP-03 | ARG | Sulfonamide | protection | CACGGCTTCGGCTCATGT | TGCCATCCTGTGACTAGCTACGT |
| dfrA1 | ARG | Sulfonamide | deactivate | GGAATGGCCCTGATATTCCA | AGTCTTGCGTCCAACCAACAG |
| dfrA12 | ARG | Sulfonamide | deactivate | CCTCTACCGAACCGTCACACA | GCGACAGCGTTGAAACAACTAC |
| folA | ARG | Sulfonamide | deactivate | CGAGCAGTTCCTGCCAAAG | CCCAGTCATCCGGTTCATAATC |
| tet(32) | ARG | Tetracycline | protection | CCATTACTTCGGACAACGGTAGA | CAATCTCTGTGAGGGCATTTAACA |
| tet(34) | ARG | Tetracycline | unknown | CTTAGCGCAAACAGCAATCAGT | CGGTGATACAGCGCGTAAACT |
| tet(35) | ARG | Tetracycline | unknown | ACCCCATGACGTACCTGTAGAGA | CAACCCACACTGGCTACCAGTT |
| tet(36)-01 | ARG | Tetracycline | protection | AGAATACTCAGCAGAGGTCAGTTCCT | TGGTAGGTCGATAACCCGAAAAT |
| tet(36)-02 | ARG | Tetracycline | protection | TGCAGGAAAGACCTCCATTACAG | CTTTGTCCACACTTCCACGTACTATG |
| tet(37) | ARG | Tetracycline | unknown | GAGAACGTTGAAAAGGTGGTGAA | AACCAAGCCTGGATCAGTCTCA |
| tetA-01 | ARG | Tetracycline | efflux | GCTGTTTGTTCTGCCGGAAA | GGTTAAGTTCCTTGAACGCAAACT |
| tetA-02 | ARG | Tetracycline | efflux | CTCACCAGCCTGACCTCGAT | CACGTTGTTATAGAAGCCGCATAG |
| tetB-01 | ARG | Tetracycline | efflux | AGTGCGCTTTGGATGCTGTA | AGCCCCAGTAGCTCCTGTGA |
| tetB-02 | ARG | Tetracycline | efflux | GCCCAGTGCTGTTGTTGTCAT | TGAAAGCAAACGGCCTAAATACA |
| tetC-01 | ARG | Tetracycline | efflux | CATATCGCAATACATGCGAAAAA | AAAGCCGCGGTAAATAGCAA |
| tetC-02 | ARG | Tetracycline | efflux | ACTGGTAAGGTAAACGCCATTGTC | ATGCATAAACCAGCCATTGAGTAAG |
| tetD-01 | ARG | Tetracycline | efflux | TGCCGCGTTTGATTACACA | CACCAGTGATCCCGGAGATAA |
| tetD-02 | ARG | Tetracycline | efflux | TGTCATCGCGCTGGTGATT | CATCCGCTTCCGGGAGAT |
| tetE | ARG | Tetracycline | efflux | TTGGCGCTGTATGCAATGAT | CGACGACCTATGCGATCTGA |
| tetG-01 | ARG | Tetracycline | efflux | TCAACCATTGCCGATTCGA | TGGCCCGGCAATCATG |
| tetG-02 | ARG | Tetracycline | efflux | CATCAGCGCCGGTCTTATG | CCCCATGTAGCCGAACCA |
| tetH | ARG | Tetracycline | efflux | TTTGGGTCATCTTACCAGCATTAA | TTGCGCATTATCATCGACAGA |
| tetJ | ARG | Tetracycline | efflux | GGGTGCCGCATTAGATTACCT | TCGTCCAATGTAGAGCATCCATA |

| | | | | | |
|---|---|---|---|---|---|
| tetK | ARG | Tetracycline | efflux | CAGCAGTCATTGGAAAATTATCTGATTATA | CCTTGTACTAACCTACCAAAAATCAAAATA |
| tetL-01 | ARG | Tetracycline | efflux | AGCCCGATTTATTCAAGGAATTG | CAAATGCTTTCCCCCTGTTCT |
| tetL-02 | ARG | Tetracycline | efflux | ATGGTTGTAGTTGCGCGCTATAT | ATCGCTGGACCGACTCCTT |
| tetM-01 | ARG | Tetracycline | protection | CATCATAGACACGCCAGGACATAT | CGCCATCTTTTGCAGAAATCA |
| tetM-02 | ARG | Tetracycline | protection | TAATATTGGAGTTTTAGCTCATGTTGATG | CCTCTCTGACGTTCTAAAAGCGTATTAT |
| tetO-01 | ARG | Tetracycline | protection | ATGTGGATACTACAACGCATGAGATT | TGCCTCCACATGATATTTTTCCT |
| tetPA | ARG | Tetracycline | efflux | AGTTGCAGATGTGTATAGTCGTAAACTATCTATT | TGCTACAAGTACGAAAACAAAACTAGAA |
| tetPB-01 | ARG | Tetracycline | protection | ACACCTGGACACGCTGATTTT | ACCGTCTAGAACGCGGAATG |
| tetPB-02 | ARG | Tetracycline | protection | TGATACACCTGGACACGCTGAT | CGTCCAAAACGCGGAATG |
| tetPB-03 | ARG | Tetracycline | protection | TGGGCGACAGTAGGCTTAGAA | TGACCCTACTGAAACATTAGAAATATACCT |
| tetPB-05 | ARG | Tetracycline | protection | CTGAAGTGGAGCGATCATTCC | CCCTCAACGGCAGAAATAACTAA |
| tetQ | ARG | Tetracycline | protection | CGCCTCAGAAGTAAGTTCATACACTAAG | TCGTTCATGCGGATATTATCAGAAT |
| tetR-02 | ARG | Tetracycline | efflux | CGCGATAGACGCCTTCGA | TCCTGACAACGAGCCTCCTT |
| tetR-03 | ARG | Tetracycline | efflux | CGCGATGGAGCAAAAGTACAT | AGTGAAAAACCTTGTTGGCATAAAA |
| tetS | ARG | Tetracycline | protection | TTAAGGACAAACTTTCTGACGACATC | TGTCTCCCATTGTTCTGGTTCA |
| tetT | ARG | Tetracycline | protection | CCATATAGAGGTTCCACCAAATCC | TGACCCTATTGGTAGTGGTTCTATTG |
| tetU-01 | ARG | Tetracycline | unknown | GTGGCAAAGCAACGGATTG | TGCGGGCTTGCAAAACTATC |
| tetV | ARG | Tetracycline | efflux | GCGGGAACGACGATGTATATC | CCGCTATCTCACGACCATGAT |
| tetW-01 | ARG | Tetracycline | protection | ATGAACATTCCCACCGTTATCTTT | ATATCGGCGGAGAGCTTATCC |
| tetX | ARG | Tetracycline | unknown | AAATTTGTTACCGACACGGAAGTT | CATAGCTGAAAAAATCCAGGACAGTT |
| vanA | ARG | Vancomycin | protection | AAAAGGCTCTGAAAACGCAGTTAT | CGGCCGTTATCTTGTAAAAACAT |
| vanB-01 | ARG | Vancomycin | protection | TTGTCGGCGAAGTGGATCA | AGCCTTTTTCCGGCTCGTT |
| vanC-01 | ARG | Vancomycin | protection | ACAGGGATTGGCTATGAACCAT | TGACTGGCGATGATTTGACTATG |
| vanC-02 | ARG | Vancomycin | protection | CCTGCCACAATCGATCGTT | CGGCTTCATTCGGCTTGATA |
| vanC-03 | ARG | Vancomycin | protection | AAATCAATACTATGCCGGGCTTT | CCGACCGCTGCCATCA |
| vanC1 | ARG | Vancomycin | protection | AGGCGATAGCGGGTATTGAA | CAATCGTCAATTGCTCATTTCC |
| vanC2/vanC3 | ARG | Vancomycin | protection | TTTGACTGTCGGTGCTTGTGA | TCAATCGTTTCAGGCAATGG |
| vanG | ARG | Vancomycin | protection | ATTTGAATTGGCAGGTATACAGGTTA | TGATTTGTCTTTGTCCATACATAATGC |
| vanHB | ARG | Vancomycin | protection | GAGGTTTCCGAGGCGACAA | CTCTCGGCGGCAGTCGTAT |
| vanHD | ARG | Vancomycin | protection | GTGGCCGATTATACCGTCATG | CGCAGGTCATTCAGGCAAT |
| vanRA-01 | ARG | Vancomycin | protection | CCCTTACTCCCACCGAGTTTT | TTCGTCGCCCCATATCTCAT |
| vanRA-02 | ARG | Vancomycin | protection | CCACTCCGGCCTTGTCATT | GCTAACCACATTCCCCTTGTTTT |
| vanRB | ARG | Vancomycin | protection | GCCCTGTCGGATGACGAA | TTACATAGTCGTCTGCCTCTGCAT |
| vanRC | ARG | Vancomycin | protection | TGCGGGAAAAACTGAACGA | CCCCCCATACGGTTTTGATTA |
| vanRC4 | ARG | Vancomycin | protection | AGTGCTTTGGCTTATCTCGAAAA | TCCGGCAGCATCACATCTAA |
| vanRD | ARG | Vancomycin | protection | TTATAATGGCAAGGATGCACTAAAGT | CGTCTACATCCGGAAGCATGA |
| vanSA | ARG | Vancomycin | protection | CGCGTCATGCTTTCAAAATTC | TCCGCAGAAAGCTCAATTTGTT |

| | | | | | |
|---|---|---|---|---|---|
| vanSB | ARG | Vancomycin | protection | GCGCGGCAAATGACAAC | TTTGCCATTTTATTCGCACTGT |
| vanSC-01 | ARG | Vancomycin | protection | ATCAACTGCGGGAGAAAAGTCT | TCCGCTGTTCCGCTTCTT |
| vanSC-02 | ARG | Vancomycin | protection | GCCATCAGCGAGTCTGATGA | CAGCTGGGATCGTTTTTCCTT |
| vanTC-01 | ARG | Vancomycin | protection | CACACGCATTTTTTCCCATCTAG | CAGCCAACAGATCATCAAAACAA |
| vanTC-02 | ARG | Vancomycin | protection | ACAGTTGCCGCTGGTGAAG | CGTGGCTGGTCGATCAAAA |
| vanTE | ARG | Vancomycin | protection | GTGGTGCCAAGGAAGTTGCT | CGTAGCCACCGCAAAAAAAT |
| vanTG | ARG | Vancomycin | protection | CGTGTAGCCGTTCCGTTCTT | CGGCATTACAGGTATATCTGGAAA |
| vanWB | ARG | Vancomycin | protection | CGGACAAAGATACCCCCTATAAAG | AAATAGTAAATTGCTCATCTGGCACAT |
| vanWG | ARG | Vancomycin | protection | ACATTTTCATTTTGGCAGCTTGTAC | CCGCCATAAGAGCCTACAATCT |
| vanXA | ARG | Vancomycin | protection | CGCTAAATATGCCACTTGGGATA | TCAAAAGCGATTCAGCCAACT |
| vanXB | ARG | Vancomycin | protection | AGGCACAAAATCGAAGATGCTT | GGGTATGGCTCATCAATCAACTT |
| vanXD | ARG | Vancomycin | protection | TAAACCGTGTTATGGGAACGAA | GCGATAGCCGTCCCATAAGA |
| vanYB | ARG | Vancomycin | protection | GGCTAAAGCGGAAGCAGAAA | GATATCCACAGCAAGACCAAGCT |
| vanYD-01 | ARG | Vancomycin | protection | AAGGCGATACCCTGACTGTCA | ATTGCCGGACGGAAGCA |
| vanYD-02 | ARG | Vancomycin | protection | CAAACGGAAGAGAGGTCACTTACA | CGGACGGTAATAGGGACTGTTC |
| acrA-01 | ARG | Multidrug | efflux | CAACGATCGGACGGGTTTC | TGGCGATGCCACCGTACT |
| acrA-02 | ARG | Multidrug | efflux | GGTCTATCACCCTACGCGCTATC | GCGCGCACGAACATACC |
| acrA-03 | ARG | Multidrug | efflux | CAGACCCGCATCGCATATT | CGACAATTTCGCGCTCATG |
| acrA-04 | ARG | Multidrug | efflux | TACTTTGCGCGCCATCTTC | CGTGCGCGAACGAACAT |
| acrB-01 | ARG | Multidrug | efflux | AGTCGGTGTTCGCCGTTAAC | CAAGGAAACGAACGCAATACC |
| acrR-01 | ARG | Multidrug | efflux | GCGCTGGAGACACGACAAC | GCCTTGCTGCGAGAACAAA |
| acrR-02 | ARG | Multidrug | efflux | GATGATACCCCCTGCTGTGAGA | ACCAAACAAGAAGCGCAAGAA |
| adeA | ARG | Multidrug | efflux | CAGTTCGAGCGCCTATTTCTG | CGCCCTGACCGACCAAT |
| acrA-05 | ARG | Multidrug | efflux | CGTGCGCGAACGAACA | ACTTTGCGCGCCATCTTC |
| acrF | ARG | Multidrug | efflux | GCGGCCAGGCACAAAA | TACGCTCTTCCCACGGTTTC |
| ceoA | ARG | Multidrug | efflux | ATCAACACGGACCAGGACAAG | GGAAAGTCCGCTCACGATGA |
| cmeA | ARG | Multidrug | efflux | GCAGCAAAGAAGAAGCACCAA | AGCAGGGTAAGTAAAACTAAGTGGTAAATCT |
| cmr | ARG | Multidrug | efflux | CGGCATCGTCAGTGGAATT | CGGTTCCGAAAAAGATGGAA |
| emrD | ARG | Multidrug | efflux | CTCAGCAGTATGGTGGTAAGCATT | ACCAGGCGCCGAAGAAC |
| marR-01 | ARG | Multidrug | efflux | GCGGCGTACTGGTGAAGCTA | TGCCCTGGTCGTTGATGA |
| mdetl1 | ARG | Multidrug | efflux | ATACAGCAGTGGATATTGGTTTAATTGT | TGCATAAGGTGAATGTTCCATGA |
| mdtE/yhiU | ARG | Multidrug | efflux | CGTCGGCGCACTCGTT | TCCAGACGTTGTACGGTAACCA |
| mepA | ARG | Multidrug | efflux | ATCGGTCGCTCTTCGTTCAC | ATAAATAGGATCGAGCTGCTGGAT |
| mexA | ARG | Multidrug | efflux | AGGACAACGCTATGCAACGAA | CCGGAAAGGGCCGAAAT |
| mexD | ARG | Multidrug | efflux | TTGCCACTGGCTTTCATGAG | CACTGCGGAGAACTGTCTGTAGA |
| mexE | ARG | Multidrug | efflux | GGTCAGCACCGACAAGGTCTAC | AGCTCGACGTACTTGAGGAACAC |

| | | | | | |
|---|---|---|---|---|---|
| mexF | ARG | Multidrug | efflux | CCGCGAGAAGGCCAAGA | TTGAGTTCGGCGGTGATGA |
| mtrC-01 | ARG | Multidrug | efflux | GGACGGGAAGATGGTCCAA | CGTAGCGTTCCGGTTCGAT |
| mtrC-02 | ARG | Multidrug | efflux | CGGAGTCCATCGACCATTTG | ATCGTCGGCAAGGAGAATCA |
| mtrD-02 | ARG | Multidrug | efflux | GGTCGGCACGCTCTTGTC | TGAAGAATTTGCGCACCACTAC |
| mtrD-03 | ARG | Multidrug | efflux | CCGCCAAGCCGATATAGACA | GGCCGGGTTGCCAAA |
| oprD | ARG | Multidrug | efflux | ATGAAGTGGAGCGCCATTG | GGCCACGGCGAACTGA |
| oprJ | ARG | Multidrug | efflux | ACGAGAGTGGCGTCGACAA | AAGGCGATCTCGTTGAGGAA |
| pmrA | ARG | Multidrug | efflux | TTTGCAGGTTTTGTTCCTAATGC | GCAGAGCCTGATTTCTCCTTTG |
| putative multidrug | ARG | Multidrug | efflux | AATTTTGCCGATTATTGCTGAAA | GATTGTCATCATTCGTTTATCACCAA |
| qac | ARG | Multidrug | efflux | CAATAATAACCGAAATAATAGGGACAAGTT | AATAAGTGTTCCTAGTGTTGGCCATAG |
| qacA | ARG | Multidrug | efflux | TGGCAATAGGAGCTATGGTGTTT | AAGGTAACACTATTTTCGGTCCAAATC |
| qacA/qacB | ARG | Multidrug | efflux | TTTAGGCAGCCTCGCTTCA | CCGAATCCAAATAAAACCCAATAA |
| qacEdelta1-01 | ARG | Multidrug | efflux | TCGCAACATCCGCATTAAAA | ATGGATTTCAGAACCAGAGAAAGAAA |
| qacEdelta1-02 | ARG | Multidrug | efflux | CCCCTTCCGCCGTTGT | CGACCAGACTGCATAAGCAACA |
| qacH-01 | ARG | Multidrug | efflux | GTGGCAGCTATCGCTTGGAT | CCAACGAACGCCCACAA |
| qacH-02 | ARG | Multidrug | efflux | CATCGTGCTTGTGGCAGCTA | TGAACGCCCAGAAGTCTAGTTTT |
| rarD-02 | ARG | Multidrug | efflux | TGACGCATCGCGTGATCT | AAATTTTCTGTGGCGTCTGAATC |
| sdeB | ARG | Multidrug | efflux | CACTACCGCTTCCGCACTTAA | TGAAAAAACGGGAAAAGTCCAT |
| tolC-01 | ARG | Multidrug | efflux | GGCCGAGAACCTGATGCA | AGACTTACGCAATTCCGGGTTA |
| tolC-02 | ARG | Multidrug | efflux | CAGGCAGAGAACCTGATGCA | CGCAATTCCGGGTTGCT |
| tolC-03 | ARG | Multidrug | efflux | GCCAGGCAGAGAACCTGATG | CGCAATTCCGGGTTGCT |
| ttgA | ARG | Multidrug | efflux | ACGCCAATGCCAAACGATT | GTCACGGCGCAGCTTGA |
| ttgB | ARG | Multidrug | efflux | TCGCCCTGGATGTACACCTT | ACCATTGCCGACATCAACAAC |
| yceE/mdtG-01 | ARG | Multidrug | efflux | TGGCACAAAATATCTGGCAGTT | TTGTGTGGCGATAAGAGCATTAG |
| yceE/mdtG-02 | ARG | Multidrug | efflux | TTATCTGTTTTCTGCTCACCTTCTTTT | GCGTGGTGACAAACAGGCTTA |
| yceL/mdtH-01 | ARG | Multidrug | efflux | TCGGGATGGTGGGCAAT | CGATAACCGAGCCGATGTAGA |
| yceL/mdtH-02 | ARG | Multidrug | efflux | CGCGTGAAACCTTAAGTGCTT | AGACGGCTAAACCCCATATAGCT |
| yceL/mdtH-03 | ARG | Multidrug | efflux | CTGCCGTTAAATGGATGTATGC | ACTCCAGCGGGCGATAGG |
| yidY/mdtL-01 | ARG | Multidrug | efflux | GCAGTTGCATATCGCCTTCTC | CTTCCCGGCAAACAGCAT |
| yidY/mdtL-02 | ARG | Multidrug | efflux | TGCTGATCGGGATTCTGATTG | CAGGCGCGACGAACATAAT |
| fabK | ARG | other | deactivate | TTTCAGCTCAGCACTTTGGTCAT | AAGGCATCTTTTTCAGCCAGTTC |
| imiR | ARG | other | unknown | CCGGACTAGAGCTTCATGTAAGC | CCCACGCGGTACTCTTGTAAA |
| nisB | ARG | other | unknown | GGGAGAGTTGCCGATGTTGTA | AGCCACTCGTTAAAGGGCAAT |
| speA | ARG | other | unknown | GCAAGAGGTATTTGCTCAACAAGA | CAGGGTCACCCTCATAAAGAAAA |
| bacA-01 | ARG | other | deactivate | CGGCTTCGTGACCTCGTT | ACAATGCGATACCAGGCAAAT |
| bacA-02 | ARG | other | deactivate | TTCCACGACACGATTAAGTCATTG | CGGCTCTTTCGGCTTCAG |

| | | | | | |
|---|---|---|---|---|---|
| fosB | ARG | other | deactivate | TCACTGTAACTAATGAAGCATTAGACCAT | CCATCTGGATCTGTAAAGTAAAGAGATC |
| fosX | ARG | other | deactivate | GATTAAGCCATATCACTTTAATTGTGAAAG | TCTCCTTCCATAATGCAAATCCA |
| nimE | ARG | other | unknown | TGCGCCAAGATAGGGCATA | GTCGTGAATTCGGCAGGTTTA |
| pncA | ARG | other | unknown | GCAATCGAGGCGGTGTTC | TTGCCGCAGCCAATTCA |
| sat4 | ARG | other | deactivate | GAATGGGCAAAGCATAAAAACTTG | CCGATTTTGAAACCACAATTATGATA |

**Supplementary Table 2.** Biomes included in this study. The biome classification was done based on vegetation field information and climatic information from the Köppen classification[16].

| Ecosystem | n | Details |
|---|---|---|
| Boreal Forests | 64 | Northern Hemisphere, high latitude Forests (>50º) |
| Cold Forests | 87 | Continental Forests excluding boreal Forests |
| Temp. Forests | 260 | Temperate Forests (67% Subtropical, 19% Oceanic and 14% Mediterranean) |
| Trop. Forests | 24 | Tropical Forests |
| Dry Forests | 105 | Arid Forests |
| Cold Grasslands | 19 | Continental Grasslands |
| Temp./trop. Grasslands | 28 | Temperate and tropical Grasslands |
| Dry Grasslands | 47 | Arid Grasslands |
| Cold Shrublands | 3 | Continental Shrublands |
| Temp./trop. Shrublands | 30 | Temperate and tropical Shrublands |
| Dry Shrublands | 122 | Arid Shrublands |
| Tundra | 13 | Polar climate |
| Croplands | 210 | Croplands including maize, soybean, rice and peanuts |

**Supplementary Table 3.** Environmental factors included in our structural equation model.

| Variables | Label | Variable | Units |
|---|---|---|---|
| ARGs | ARGs | Richness or proportion | Number of ARGs / Standardized relative abundance of ARGs |
| Spatial | Elevation | Elevation | m |
| | Spatial dissimilarity | (Euclidean distance) based on latitude and longitude (sine and cosine) | Decimal degrees |
| Climate | MAT | Mean annual temperature | ºC |
| | MAP | Mean annual precipitation | mm |
| | TSEA | Temperature Seasonality (standard deviation *100) | Unitless |
| | PSEA | Precipitation Seasonality (Coefficient of Variation) | % |
| Vegetation | Forests (F) | Location supporting Forests ecosystems | 0/1 |
| | Grasslands (G) | Location supporting Grasslands ecosystems | 0/1 |
| | Shrublands (S) | Location supporting Shrublands ecosystems | 0/1 |
| | Plant cover | Plant cover (vegetation cover) | % * 100 |
| Soil | Fine texture (Clay+silt) | Fine texture | % |
| | pH | pH | Unitless |
| | Soil C | Soil carbon | % |
| | Soil C:N | Soil carbon to nitrogen ratio | Unitless |
| MGEs | MGEs | Richness and proportion | Number of MGEs / Standardized MGE relative abundance |

**Supplementary Table 4.** Standardized direct effects of SEM on the proportion of soil ARGs.

| Parameters | | | Direct effect | Bootstrap-P |
|---|---|---|---|---|
| PSEA | ← | Spatial dissimilarity | -0.176 | 0.001 |
| MAT | ← | Spatial dissimilarity | -0.028 | 0.264 |
| MAP | ← | Spatial dissimilarity | -0.322 | 0.001 |
| TSEA | ← | Spatial dissimilarity | -0.292 | 0.001 |
| PSEA | ← | Elevation | 0.218 | 0.001 |
| MAT | ← | Elevation | -0.200 | 0.001 |
| MAP | ← | Elevation | -0.199 | 0.001 |
| TSEA | ← | Elevation | -0.136 | 0.001 |
| Forests | ← | PSEA | 0.114 | 0.001 |
| Forests | ← | MAT | -0.088 | 0.271 |
| Forests | ← | MAP | 0.343 | 0.001 |
| Forests | ← | TSEA | 0.029 | 0.715 |
| Plant cover | ← | PSEA | 0.114 | 0.001 |
| Plant cover | ← | MAT | -0.502 | 0.002 |
| Plant cover | ← | MAP | 0.677 | 0.001 |
| Plant cover | ← | TSEA | -0.513 | 0.002 |
| Grasslands | ← | PSEA | -0.072 | 0.090 |
| Shrublands | ← | PSEA | -0.052 | 0.190 |
| Grasslands | ← | MAT | -0.166 | 0.026 |
| Shrublands | ← | MAT | 0.110 | 0.109 |
| Grasslands | ← | MAP | -0.231 | 0.001 |
| Shrublands | ← | MAP | -0.509 | 0.001 |
| Grasslands | ← | TSEA | -0.274 | 0.001 |
| Shrublands | ← | TSEA | -0.144 | 0.101 |
| Plant cover | ← | Spatial dissimilarity | -0.261 | 0.001 |
| Forests | ← | Spatial dissimilarity | 0.245 | 0.002 |
| Shrublands | ← | Spatial dissimilarity | -0.154 | 0.005 |
| Grasslands | ← | Spatial dissimilarity | 0.021 | 0.680 |
| Plant cover | ← | Elevation | -0.321 | 0.001 |
| Forests | ← | Elevation | 0.015 | 0.733 |
| Shrublands | ← | Elevation | 0.120 | 0.012 |
| Grasslands | ← | Elevation | 0.105 | 0.051 |
| Soil C:N | ← | Forests | 0.503 | 0.002 |
| Soil C | ← | Forests | 0.342 | 0.001 |
| pH | ← | Forests | -0.418 | 0.002 |
| pH | ← | Plant cover | 0.041 | 0.423 |
| Soil C | ← | Plant cover | 0.008 | 0.895 |

| | | | | |
|---|---|---|---|---|
| Soil C:N | ← | Plant cover | -0.037 | 0.527 |
| pH | ← | PSEA | -0.038 | 0.251 |
| Soil C | ← | PSEA | 0.016 | 0.511 |
| Soil C:N | ← | PSEA | -0.099 | 0.009 |
| pH | ← | MAT | 0.227 | 0.001 |
| Soil C | ← | MAT | -0.803 | 0.002 |
| pH | ← | MAP | -0.667 | 0.001 |
| Soil C | ← | MAP | 0.013 | 0.747 |
| Soil C:N | ← | MAP | 0.141 | 0.025 |
| pH | ← | TSEA | -0.034 | 0.568 |
| Soil C | ← | TSEA | -0.513 | 0.001 |
| Soil C:N | ← | TSEA | -0.314 | 0.001 |
| Soil C:N | ← | Shrublands | 0.272 | 0.002 |
| Soil C | ← | Shrublands | 0.002 | 0.936 |
| pH | ← | Shrublands | -0.111 | 0.002 |
| Soil C:N | ← | Grasslands | 0.135 | 0.010 |
| Soil C | ← | Grasslands | -0.005 | 0.900 |
| pH | ← | Grasslands | -0.174 | 0.002 |
| Soil C | ← | Spatial dissimilarity | -0.251 | 0.001 |
| Soil C:N | ← | Spatial dissimilarity | 0.004 | 0.937 |
| pH | ← | Spatial dissimilarity | -0.077 | 0.023 |
| Fine texture | ← | Plant cover | 0.049 | 0.298 |
| Fine texture | ← | Forests | -0.376 | 0.001 |
| Fine texture | ← | Shrublands | -0.377 | 0.001 |
| Fine texture | ← | Grasslands | -0.331 | 0.001 |
| Fine texture | ← | PSEA | 0.082 | 0.006 |
| Fine texture | ← | MAP | 0.052 | 0.238 |
| Fine texture | ← | TSEA | 0.249 | 0.002 |
| Fine texture | ← | Spatial dissimilarity | -0.143 | 0.001 |
| Soil C:N | ← | MAT | -0.513 | 0.001 |
| Fine texture | ← | Elevation | -0.114 | 0.004 |
| pH | ← | Elevation | 0.153 | 0.001 |
| Soil C | ← | Elevation | -0.090 | 0.148 |
| Soil C:N | ← | Elevation | -0.074 | 0.098 |
| Fine texture | ← | MAT | -0.150 | 0.031 |
| MGE proportion | ← | Plant cover | 0.052 | 0.420 |
| MGE proportion | ← | Forests | 0.033 | 0.597 |
| MGE proportion | ← | Shrublands | 0.144 | 0.006 |
| MGE proportion | ← | Grasslands | -0.007 | 0.796 |
| MGE proportion | ← | Soil C:N | 0.079 | 0.139 |
| MGE proportion | ← | Soil C | 0.007 | 0.835 |

| | | | | |
|---|---|---|---|---|
| MGE proportion | ← | pH | 0.022 | 0.615 |
| MGE proportion | ← | Spatial dissimilarity | -0.045 | 0.408 |
| MGE proportion | ← | PSEA | 0.060 | 0.434 |
| MGE proportion | ← | MAT | -0.220 | 0.143 |
| MGE proportion | ← | MAP | 0.077 | 0.327 |
| MGE proportion | ← | TSEA | -0.125 | 0.490 |
| MGE richness | ← | Soil C | -0.011 | 0.777 |
| MGE richness | ← | Soil C:N | 0.032 | 0.626 |
| MGE richness | ← | pH | 0.130 | 0.003 |
| MGE richness | ← | TSEA | 0.199 | 0.018 |
| MGE richness | ← | MAP | 0.008 | 0.933 |
| MGE richness | ← | MAT | -0.053 | 0.495 |
| MGE richness | ← | PSEA | 0.133 | 0.002 |
| MGE richness | ← | Spatial dissimilarity | -0.008 | 0.822 |
| MGE richness | ← | Grasslands | -0.024 | 0.627 |
| MGE richness | ← | Shrublands | -0.055 | 0.272 |
| MGE richness | ← | Forests | -0.017 | 0.875 |
| MGE richness | ← | Plant cover | 0.255 | 0.001 |
| MGE richness | ← | Fine texture | -0.067 | 0.083 |
| MGE proportion | ← | Fine texture | -0.152 | 0.009 |
| MGE proportion | ← | Elevation | -0.058 | 0.473 |
| MGE richness | ← | Elevation | 0.063 | 0.213 |
| ARG proportion | ← | Plant cover | -0.053 | 0.169 |
| ARG proportion | ← | PSEA | 0.111 | 0.001 |
| ARG proportion | ← | MAT | -0.245 | 0.003 |
| ARG proportion | ← | MAP | 0.074 | 0.107 |
| ARG proportion | ← | TSEA | -0.183 | 0.032 |
| ARG proportion | ← | Forests | 0.078 | 0.014 |
| ARG proportion | ← | Soil C:N | -0.039 | 0.130 |
| ARG proportion | ← | Soil C | 0.003 | 0.916 |
| ARG proportion | ← | pH | 0.007 | 0.777 |
| ARG proportion | ← | Shrublands | 0.092 | 0.044 |
| ARG proportion | ← | Grasslands | 0.023 | 0.430 |
| ARG proportion | ← | Spatial dissimilarity | -0.058 | 0.241 |
| ARG proportion | ← | MGE proportion | 0.694 | 0.001 |
| ARG proportion | ← | MGE richness | -0.048 | 0.069 |
| ARG proportion | ← | Fine texture | -0.029 | 0.486 |
| ARG proportion | ← | Elevation | -0.085 | 0.050 |

**Supplementary Table 5.** Standardized direct effects of SEM on the richness of soil ARGs.

| Parameters | | | Direct effect | Bootstrap-P |
|---|---|---|---|---|
| PSEA | ← | Spatial dissimilarity | -0.176 | 0.001 |
| MAT | ← | Spatial dissimilarity | -0.028 | 0.264 |
| MAP | ← | Spatial dissimilarity | -0.322 | 0.001 |
| TSEA | ← | Spatial dissimilarity | -0.292 | 0.001 |
| PSEA | ← | Elevation | 0.218 | 0.001 |
| MAT | ← | Elevation | -0.200 | 0.001 |
| MAP | ← | Elevation | -0.199 | 0.001 |
| TSEA | ← | Elevation | -0.136 | 0.001 |
| Forests | ← | PSEA | 0.114 | 0.001 |
| Forests | ← | MAT | -0.088 | 0.271 |
| Forests | ← | MAP | 0.343 | 0.001 |
| Forests | ← | TSEA | 0.029 | 0.715 |
| Plant cover | ← | PSEA | 0.114 | 0.001 |
| Plant cover | ← | MAT | -0.502 | 0.002 |
| Plant cover | ← | MAP | 0.677 | 0.001 |
| Plant cover | ← | TSEA | -0.513 | 0.002 |
| Grasslands | ← | PSEA | -0.072 | 0.090 |
| Shrublands | ← | PSEA | -0.052 | 0.190 |
| Grasslands | ← | MAT | -0.166 | 0.026 |
| Shrublands | ← | MAT | 0.110 | 0.109 |
| Grasslands | ← | MAP | -0.231 | 0.001 |
| Shrublands | ← | MAP | -0.509 | 0.001 |
| Grasslands | ← | TSEA | -0.274 | 0.001 |
| Shrublands | ← | TSEA | -0.144 | 0.101 |
| Plant cover | ← | Spatial dissimilarity | -0.261 | 0.001 |
| Forests | ← | Spatial dissimilarity | 0.245 | 0.002 |
| Shrublands | ← | Spatial dissimilarity | -0.154 | 0.005 |
| Grasslands | ← | Spatial dissimilarity | 0.021 | 0.680 |
| Plant cover | ← | Elevation | -0.321 | 0.001 |
| Forests | ← | Elevation | 0.015 | 0.733 |
| Shrublands | ← | Elevation | 0.120 | 0.012 |
| Grasslands | ← | Elevation | 0.105 | 0.051 |
| Soil C:N | ← | Forests | 0.503 | 0.002 |
| Soil C | ← | Forests | 0.342 | 0.001 |
| pH | ← | Forests | -0.418 | 0.002 |
| pH | ← | Plant cover | 0.041 | 0.423 |
| Soil C | ← | Plant cover | 0.008 | 0.895 |

| | | | | |
|---|---|---|---|---|
| Soil C:N | ← | Plant cover | -0.037 | 0.527 |
| pH | ← | PSEA | -0.038 | 0.251 |
| Soil C | ← | PSEA | 0.016 | 0.511 |
| Soil C:N | ← | PSEA | -0.099 | 0.009 |
| pH | ← | MAT | 0.227 | 0.001 |
| Soil C | ← | MAT | -0.803 | 0.002 |
| pH | ← | MAP | -0.667 | 0.001 |
| Soil C | ← | MAP | 0.013 | 0.747 |
| Soil C:N | ← | MAP | 0.141 | 0.025 |
| pH | ← | TSEA | -0.034 | 0.568 |
| Soil C | ← | TSEA | -0.513 | 0.001 |
| Soil C:N | ← | TSEA | -0.314 | 0.001 |
| Soil C:N | ← | Shrublands | 0.272 | 0.002 |
| Soil C | ← | Shrublands | 0.002 | 0.936 |
| pH | ← | Shrublands | -0.111 | 0.002 |
| Soil C:N | ← | Grasslands | 0.135 | 0.010 |
| Soil C | ← | Grasslands | -0.005 | 0.900 |
| pH | ← | Grasslands | -0.174 | 0.002 |
| Soil C | ← | Spatial dissimilarity | -0.251 | 0.001 |
| Soil C:N | ← | Spatial dissimilarity | 0.004 | 0.937 |
| pH | ← | Spatial dissimilarity | -0.077 | 0.023 |
| Fine texture | ← | Plant cover | 0.049 | 0.298 |
| Fine texture | ← | Forests | -0.376 | 0.001 |
| Fine texture | ← | Shrublands | -0.377 | 0.001 |
| Fine texture | ← | Grasslands | -0.331 | 0.001 |
| Fine texture | ← | PSEA | 0.082 | 0.006 |
| Fine texture | ← | MAP | 0.052 | 0.238 |
| Fine texture | ← | TSEA | 0.249 | 0.002 |
| Fine texture | ← | Spatial dissimilarity | -0.143 | 0.001 |
| Soil C:N | ← | MAT | -0.513 | 0.001 |
| Fine texture | ← | Elevation | -0.114 | 0.004 |
| pH | ← | Elevation | 0.153 | 0.001 |
| Soil C | ← | Elevation | -0.090 | 0.148 |
| Soil C:N | ← | Elevation | -0.074 | 0.098 |
| Fine texture | ← | MAT | -0.150 | 0.031 |
| MGE proportion | ← | Plant cover | 0.052 | 0.420 |
| MGE proportion | ← | Forests | 0.033 | 0.597 |
| MGE proportion | ← | Shrublands | 0.144 | 0.006 |
| MGE proportion | ← | Grasslands | -0.007 | 0.796 |
| MGE proportion | ← | Soil C:N | 0.079 | 0.139 |
| MGE proportion | ← | Soil C | 0.007 | 0.835 |

| | | | | |
|---|---|---|---|---|
| MGE proportion | ← | pH | 0.022 | 0.615 |
| MGE proportion | ← | Spatial dissimilarity | -0.045 | 0.408 |
| MGE proportion | ← | PSEA | 0.060 | 0.434 |
| MGE proportion | ← | MAT | -0.220 | 0.143 |
| MGE proportion | ← | MAP | 0.077 | 0.327 |
| MGE proportion | ← | TSEA | -0.125 | 0.490 |
| MGE richness | ← | Soil C | -0.011 | 0.777 |
| MGE richness | ← | Soil C:N | 0.032 | 0.626 |
| MGE richness | ← | pH | 0.130 | 0.003 |
| MGE richness | ← | TSEA | 0.199 | 0.018 |
| MGE richness | ← | MAP | 0.008 | 0.933 |
| MGE richness | ← | MAT | -0.053 | 0.495 |
| MGE richness | ← | PSEA | 0.133 | 0.002 |
| MGE richness | ← | Spatial dissimilarity | -0.008 | 0.822 |
| MGE richness | ← | Grasslands | -0.024 | 0.627 |
| MGE richness | ← | Shrublands | -0.055 | 0.272 |
| MGE richness | ← | Forests | -0.017 | 0.875 |
| MGE richness | ← | Plant cover | 0.255 | 0.001 |
| MGE richness | ← | Fine texture | -0.067 | 0.083 |
| MGE proportion | ← | Fine texture | -0.152 | 0.009 |
| MGE proportion | ← | Elevation | -0.058 | 0.473 |
| MGE richness | ← | Elevation | 0.063 | 0.213 |
| ARG richness | ← | Plant cover | 0.237 | 0.001 |
| ARG richness | ← | PSEA | 0.054 | 0.065 |
| ARG richness | ← | MAT | 0.106 | 0.107 |
| ARG richness | ← | MAP | -0.010 | 0.812 |
| ARG richness | ← | TSEA | 0.309 | 0.001 |
| ARG richness | ← | Forests | 0.335 | 0.001 |
| ARG richness | ← | Soil C:N | 0.046 | 0.154 |
| ARG richness | ← | Soil C | -0.020 | 0.523 |
| ARG richness | ← | pH | -0.027 | 0.436 |
| ARG richness | ← | Shrublands | 0.225 | 0.001 |
| ARG richness | ← | Grasslands | 0.158 | 0.001 |
| ARG richness | ← | Spatial dissimilarity | -0.048 | 0.213 |
| ARG richness | ← | MGE proportion | 0.003 | 0.907 |
| ARG richness | ← | MGE richness | 0.476 | 0.001 |
| ARG richness | ← | Fine texture | -0.003 | 0.959 |
| ARG richness | ← | Elevation | 0.048 | 0.179 |
